# A principal-stress rule for cell division in epithelia

**DOI:** 10.64898/2026.07.15.738393

**Authors:** Lucas Anger, Tianxiang Ma, Fanny Wodrascka, Andreas Schoenit, Kristian Thijssen, Satish Kailasam Mani, Marc-Antoine Fardin, Carine Rosse, René-Marc Mège, Amin Doostmohammadi, Benoit Ladoux

## Abstract

Dense active materials, from cellular tissues to jammed and glassy systems, must continuously relieve internal mechanical stress to remain structurally stable as they are driven far from equilibrium. In epithelial tissues, this relief occurs through cell division, yet what sets the geometry of this structural remodeling event has, for over a century, been attributed to a purely geometric principle: Hertwig’s rule, whereby cells divide along their long axis. We show that as epithelial tissues densify and cell shape anisotropy collapses, this geometric rule is superseded by a mechanical one in which dense epithelia relieve anisotropic stress by cells dividing along their principal axis, independent of the tissue’s isotropic stress state. Using direct force measurement and stress inference, we show that stress orientation, rather than cell shape, governs the axis of cell division across mechanically distinct systems, from fluid-like to jammed monolayers and structurally heterogeneous organoids, remaining predictive precisely where the classical geometric rule fails. This stress-oriented remodeling is reciprocally coupled to the material’s mechanical state: anisotropic stress accelerates the underlying remodeling rate, while each remodeling event locally dissipates the stress that triggered it, closing a negative feedback loop. This “principal-stress rule” recasts epithelial cell division as a stress-relief mechanism intrinsic to dense active matter, providing a general mechanical framework linking internal stress, structural remodeling, and homeostasis in living materials.

## Main text

Cell division is a central process in development, homeostasis, and disease, and its regulation has long been studied from mechanical perspectives. Classic experiments dating back to the 19th century led to Hertwig’s rule, which states that cells orient their division along their long axis, reflecting the geometry of interphase shape [1, 2]. This geometric principle has since been refined in diverse contexts, including epithelial layers where cell shape and mitotic spindle alignment appear coupled [3–6].

Beyond geometry, accumulating evidence points to a central role for forces and stresses. Tension has been shown to accelerate proliferation [7–9], compression can inhibit it [10–12], and external forces can orient spindles independently of cell shape [13]. Within tissues, cells have also been observed to divide under compression or in mechanically ambiguous states, raising questions about whether shape or stress provides the dominant physical signal [14–20]. Despite extensive research, a unified framework reconciling geometry and stress-based control of cell division has remained elusive.

Dense epithelial tissues provide a particularly stringent test. As cells proliferate and crowd, they round up and adopt isotropic shapes [21–23]. In this regime, orientational order defined by cell shape or actin organization collapses, leaving Hertwig’s geometric rule no longer a reliable predictor. Yet tissue-level regulation of divisions must continue [24, 25], suggesting that a more general principle must govern their orientation.

Here we show that in dense epithelia, division is governed not by geometry but by mechanics. Cells align their division axis with the direction of maximal anisotropic stress, a “principal-stress rule” that generalizes Hertwig’s principle to tissues where shape cues fail, while remaining equally predictive when such cues are present. Notably, this principle is independent of the isotropic stress state, as mechanical anisotropy can exist even in the absence of a net isotropic stress. We find across different cell types and varying cell densities that stress orientation robustly governs division orientation, even as tissues jam and cell elongation disappears. Moreover, anisotropic stress not only predicts division orientation but also accelerates the cell cycle, while cytokinesis in turn dissipates anisotropy. This establishes a feedback loop that links stress, division, and homeostasis within a unified mechanical framework.

### A principal-stress rule extends Hertwig’s long-axis rule

For over a century, Hertwig’s long-axis rule has been the dominant paradigm for understanding division orientation [1, 2]. We first revisited this rule in various types of epithelial monolayers including cell lines and intestinal organoids. Starting with Madin-Darby Canine Kidney (MDCK) cells, we observed that at low density cells were elongated (Fig. 1a–b) and divisions aligned with the long axis, consistent with both the classical rule and modern refinements linking spindle positioning to geometry [3, 4, 6] (Fig. 1c–d). This served as an internal validation that our system recapitulates established principles.

**Fig. 1.**
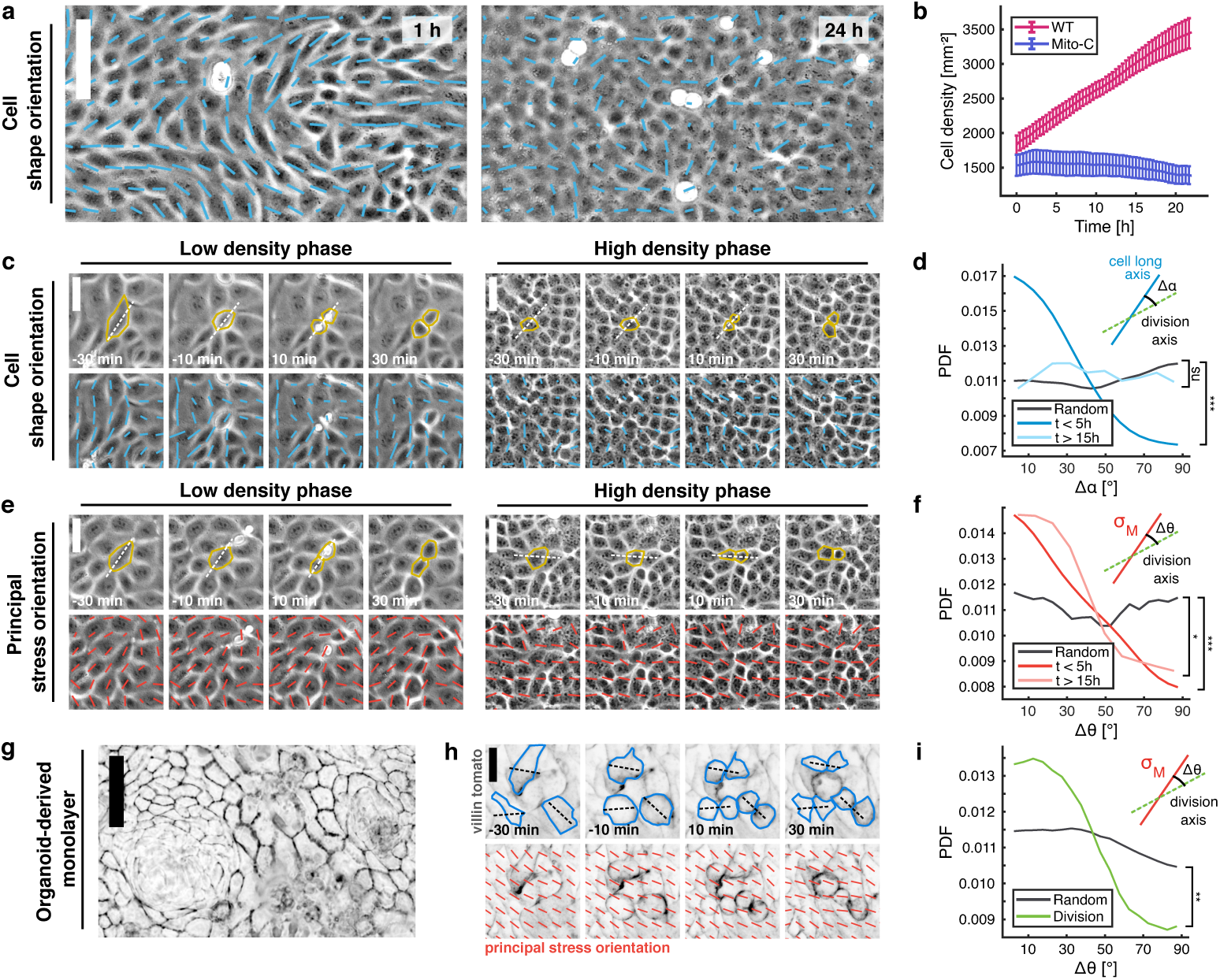
Principal stress orientation predicts cell division orientation throughout tissue densification. **a**) Left, phase contrast image of confluent MDCK monolayer in low density state (t = 1h), right, same monolayer in high density state (t = 24h). Nematic director field in blue. Magnitude of the elongation is weighted by the aspect ratio. **b**) Average (mean) evolution in time of the cell density within the whole tissue, for MDCK WT and for mitomycin-C treated MDCK cells. n = 15 movies from N = 3 independent experiments (WT); n = 15 movies from N = 3 independent experiments (Mito-C). **c**) Phase contrast image of cell division process, showing in dotted white line cell division orientation, before and after cytokinesis for (left) a low density MDCK monolayer with nematic behavior and for (right), the same monolayer at high density. Nematic director field in blue, weighted by the local aspect ratio. **d**) Probability distribution function of the angle between MDCK cell division orientation and nematic director orientation for different cell density. 681 cell divisions (t *<* 5h), 234 cell divisions (t *>* 15h), 1690 random orientations at random positions (compared with the local nematic director orientation). n = 13 movies from N = 3 independent experiments. p *≤* 0.001 (t *<* 5h vs random) and p = 0.163 (t *>* 15h vs random). **e**) Phase contrast image of cell division process, showing in dotted white line cell division orientation, before and after cytokinesis for (left) a low density MDCK monolayer with nematic behavior and for (right), the same monolayer at high density. Principal stress orientation in red. **f**) Probability distribution function of the angle between MDCK cell division orientation and principal stress orientation for different cell density. 681 cells divisions (t *<* 5h), 234 cells divisions (t *>* 15h), 1690 random orientation at random position (compared with the local principal stress orientation). n = 13 movies from N = 3 independent experiments. p *≤* 0.001 (t *<* 5h vs random) and p *≤* 0.03(t *>* 15h vs random). **g**) Phase contrast image of 2D confluent intestinal organoid-derived monolayer after being cultivated for three days. **h**) Confocal images of cell division process, showing in dotted black line cell divisions orientations, before and after cytokinesis for villintomato intestinal organoid monolayer. Principal stress orientation in red. **i**) Probability distribution function of the angle between organoid cell division orientation and principal stress orientation. 620 cell divisions and 1950 random positions. n = 13 movies from N = 3 independent experiments. p *≤* 0.005 (stress vs random). [Scale bar 80 µm (a, g), 20 µm (c, e, h). P-value from Mann-Whitney U-test (d, f, i). All error bars show the confidence interval at 95 %.]

As confluent epithelial tissues densified, however, cell geometry changed dramatically: elongated shapes gave way to isotropic, hexagonally packed cells (Fig. S1a-d), consistent with previous studies [26, 27]. In this regime, the long axis is weakly defined (Fig. S1e-f, Movie S1), and divisions are uncoupled from cell shape, indicating that a cue other than geometry must dominate (Fig. 1c–d). The breakdown of Hertwig’s rule in this regime highlights a conceptual gap: while geometric alignment explains orientation in sparse epithelia, it fails in precisely those crowded contexts where control of division orientation is most consequential for tissue architecture [19, 24, 28].

If geometry cannot explain division orientation in dense tissues, what alternative cue does? A natural candidate for such a cue is mechanics. Cells continuously generate, transmit, and respond to forces [29, 30] that can, in principle, provide directional information. We therefore asked whether local stress fields, rather than geometry, might provide the directional information that persists when shapes become isotropic. To test whether division orientation reflects stress rather than shape, we first measured the traction forces exerted by cells on their substrates (Fig. S2a-b) and subsequently inferred the stresses using Bayesian inference stress microscopy [31, 32] (Fig. S2c-e, Movie S2). At any given point of space within the epithelial sheet, the local in-plane coordinate system can be rotated to identify the specific directions along which the normal stress reaches its maximum and minimum values [33] (Fig. S2f). These directions define the maximal principal stress, *σ_M_*, the minimal principal stress, *σ_m_* and the principal stress orientation *θ* (see Supporting Information). Physically, this manipulation corresponds to separating the stress tensor into isotropic and deviatoric components, whose invariants define the isotropic stress 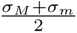 (Fig. S2c–d) and the anisotropic stress 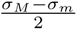, the latter being applied along the principal stress direction (Fig. S2e–g). Interestingly, temporal autocorrelation analysis indicated that principal stress orientation was slightly more temporally stable than shape anisotropy (Fig. S2h).

Across densities, division orientation was consistently aligned with the principal stress axis (Fig. 1e-f, Fig. S3a). Importantly, this “principal-stress rule” also held in intestinal epithelial organoids [34, 35], despite their heterogeneous distributions of cell types, shapes, and sizes (Fig. 1g-i, Fig. S3b, Movie S3). Finally, to test the generality of our findings, we examined epithelial cell lines covering a range of mechanical behaviors, from fluid-like mammary gland MCF10A cells (Fig. S3c–e), in which elongation persists over long times [36], to solid-like N/TERT1 keratinocytes (Fig. S3f–h) [37], which jam early and do not exhibit elongated shapes. Across these diverse mechanical contexts, including tension, compression, fluid-like and jammed regimes, and the mixed cell-type monolayers of intestinal organoids, stress orientation consistently provided a robust predictor of the division axis.

This generality is key. Prior reports noted cases where cells divided orthogonally to their shape, along an anisotropic tension or perpendicular to mechanical compression [3, 14–17, 19, 28, 38], but lacked a unifying principle. In all experimental conditions analyzed in this study, divisions are aligned with the stress axis. This principal stress orientation can emerge under compressive, tensile, or even zero isotropic stress conditions (Fig. 2a–b). In our data, tissue mechanical anisotropy is created under both compression or tension, and remained systematically non-zero during monolayer densification, thereby defining a stress orientation (Fig. 2c and Fig. S3a-b, e, h). Thus, the principal-stress rule is a general framework: even if geometric cues vanish, stress anisotropy continues to orient divisions and extends Hertwig’s rule regardless of the isotropic stress state.

**Fig. 2.**
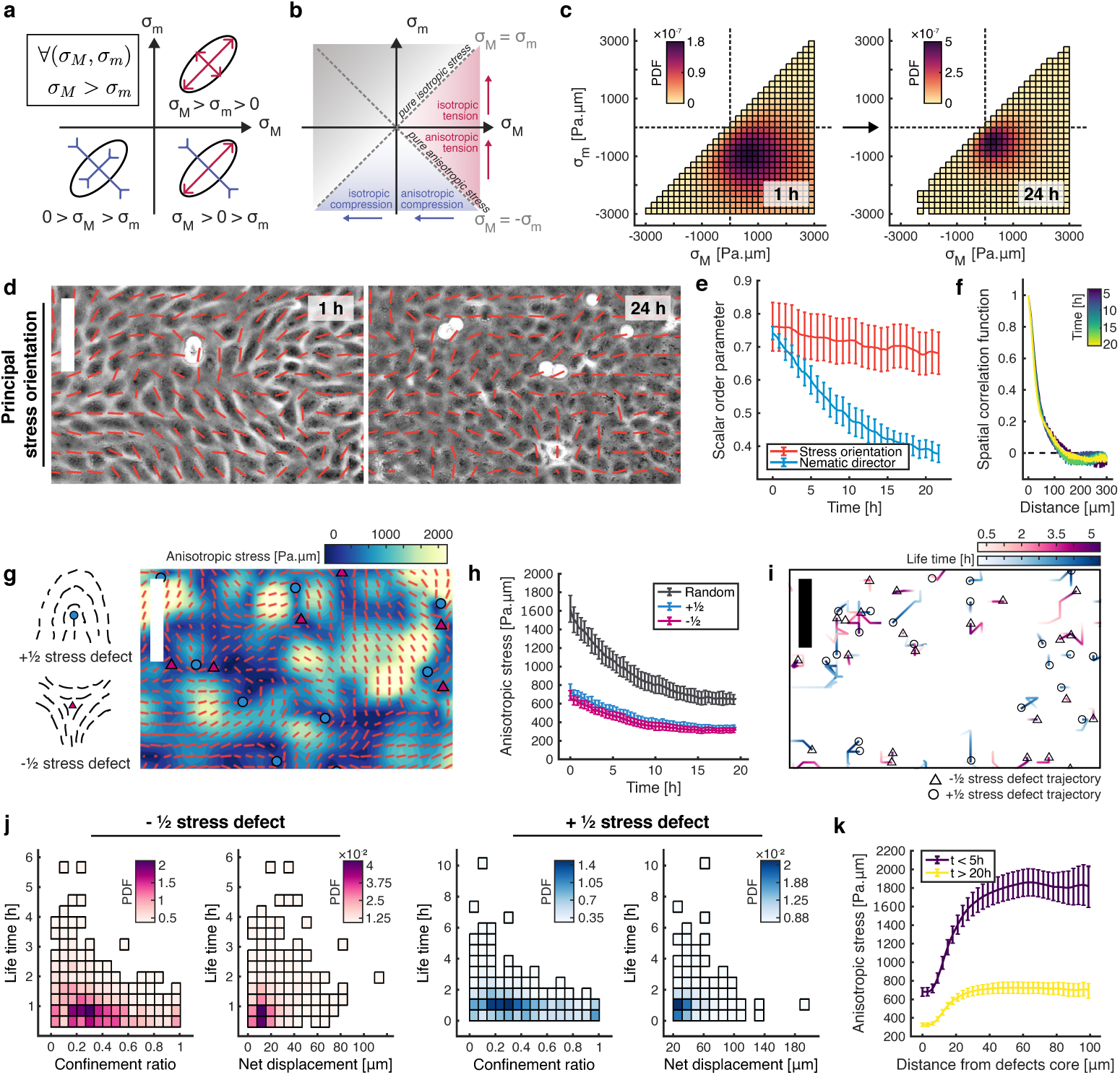
Anisotropic stress defines tissue mechanical landscape in space and in time. **a)** Mechanical state diagram illustrating all possible coupling between maximum and minimum principal stress. **b)** Mechanical state diagram illustrating all possible coupling between maximum and minimum principal stress in order to create tension or compression within the tissue. The red area corresponds to positive isotropic stress (tension), and the blue area to negative isotropic stress (compression). The gray area is prevented by construction of σ*_M_* and σ*_m_*. **c)** 2D histogram of the maximum and minimum principal stress coupling within MDCK monolayer, left, in low density state (t = 1h), and right, in high density state (t = 24h). Color code is the PDF. n = 24 movies from N = 5 independent experiments. **d)** Left, phase contrast image of low density MDCK monolayer (t = 1h) in low density state, right, same monolayer in high density state (t = 24h). Principal stress orientation field in red. Magnitude of the elongation is weighted by the anisotropic aspect ratio (see Supporting Information). **e)** Evolution of the average apparent scalar order parameter for the nematic director (blue) and for the principal stress orientation (red) in time. n = 15 movies from N = 3 independent experiments. **f)** Decay of the spatial correlation function of the principal stress orientation for various moments in time. Color code is the time. n = 15 movies from N = 3 independent experiments. **g)** Left, cartoon of various topological stress defects charge possible in epithelia, right, anisotropic stress field snapshot within the monolayer (t = 10h). Principal stress orientation in red. +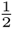 defects in blue circle, −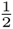 defects in pink triangle. **h)** Average anisotropic stress evolution during jamming transition around stress defects and random positions. n = 24 movies from N = 5 independent experiments. 100 random positions were taken for each frame, and on average, 50 defects of each kind were detected for one given frame. **i)** Representative stress defects trajectory within a 5h time window. +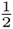 defects in blue circle, −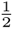 defects in pink triangle. **j)** 2D histogram of stress defects lifetime regarding, left, confinement ratio and right, net displacement. The confinement ratio ranges from 0 (confined or caged motion) to 1 (straight-line trajectory). +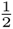 stress defects in blue (n = 2358), −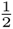 stress defects in pink (n = 1897). n = 5 movies from N = 5 independent experiments. Color code is the probability density function (PDF). **k)** Average anisotropic stress evolution regarding distance from defects core, regardless of the defect charge, during jamming transition. n = 24 movies from N = 5 independent experiments. On average, 50 defects of each kind were detected for one given frame. [Scale bar 80 μm (d, g, i). All error bars show the confidence interval at 95 %.]

### Stress orientation persists when orientational order collapses

Building on the stress metrics defined above, we next investigated how stress orientations evolve over time. Indeed, prior studies have revealed that cellular monolayers could display orientational order, such as nematic domains arising from cell shape or actin alignment [26, 39, 40], which by proxy would be expected to influence division orientation through Hertwig’s rule. We therefore compared how shape-based versus stress-based order evolve as tissues densify. At low density, MDCK monolayers exhibited nematic domains spanning hundreds of microns, with actin fibers aligned across cells (Fig. S4a). These orientational cues coincided with division alignment, consistent with the idea that collective order provides the directional bias. With increasing cell density, however, nematic order collapsed and cells adopted isotropic packing (Fig. S4a-d). This loss occurred across scales, single-cell actin ordering also disappeared at high density, confirming that orientational cues derived from geometry or cytoskeletal alignment vanish in crowded epithelia (Fig. S4d-e), without being replaced by an apico-basal anisotropy arising from mismatches between the apical and basal cell surfaces (Fig. S4f).

In contrast, stress orientation remained highly ordered. Even in jammed tissues [41], where motion slowed and shape order was lost (Fig. S5a-d), stress domains persisted with strong alignment (Fig. 2d–e). Importantly, the scale of stress domains did not shrink with density; rather, anisotropy extended over 5-6 cell diameters (Fig. 2f). This persistence explains why division orientation remains coherent even as local geometry becomes isotropic (Movie S4). To further probe the limits of stress orientation in our system and test its quantification by BISM, we imposed specific stress patterns using controlled boundary conditions via micropatterning [42]. Confining cells to elliptical patterns generated a principal stress orientation along the major axis, along which cells preferentially divided (Fig. S6a-b). Consistently, laser ablation induced recoil predominantly along the major axis (Fig. 6c-e), independently validating both the ability of epithelia to adopt a collective anisotropic stress organization and the ability of BISM to recover the orientation of the deviatoric stress tensor [32].

These results highlight an essential distinction: geometric order is instable, collapsing under crowding, whereas stress order is robust, integrating forces across many cells. Stress anisotropy therefore provides a global orientational field that is insulated from local packing fluctuations. In dense tissues, where reliable alignment is needed to prevent uncontrolled anisotropy [43], stress-based order emerges as the dominant organizing principle.

Beyond persistent order, stress fields also exhibited distinct topological defects. Unlike shape- or actin-based nematic defects, which disappeared as density increased, defects in the stress field persisted even in jammed monolayers (Fig. 2g and Fig. S7a-b) and remained spatiotemporally stable (Fig. 2h-j and Fig. S7c-f). To test whether these structures influence proliferation, we measured the spatial distribution between defects and division events. Divisions occurred at an average distance of ∼ 35*µm* from the nearest defect, nearly half the mean spacing between defects of the same type (∼ 65*µm*, Fig. S8a–c). This indicates that divisions do not occur at defect cores, but preferentially at intermediate distances where stress anisotropy is elevated (Fig. 2k and Fig. S8d-e) and stress orientation is well defined. In this sense, defects act as organizing centers that spatially repel divisions while promoting them in surrounding high-anisotropy regions.

### Anisotropic stress regulates cell cycle progression

While orientation determines *where* cells divide, proliferation also depends on *when*. Mechanical signals have long been implicated in proliferation [44], with tension reported to promote cell cycle entry [7, 8] and compression to inhibit it [10–12]. However, these models focus on isotropic stress, leaving open whether anisotropy itself plays a role. In our experiments, nearly 40% of divisions occurred in compressed or isotropically neutral regions (Fig. 3a), inconsistent with a purely tension-based explanation. Instead, it was high anisotropy, the directional imbalance of stress, that correlated with more cytokinesis events (Fig. 3b-c).

**Fig. 3.**
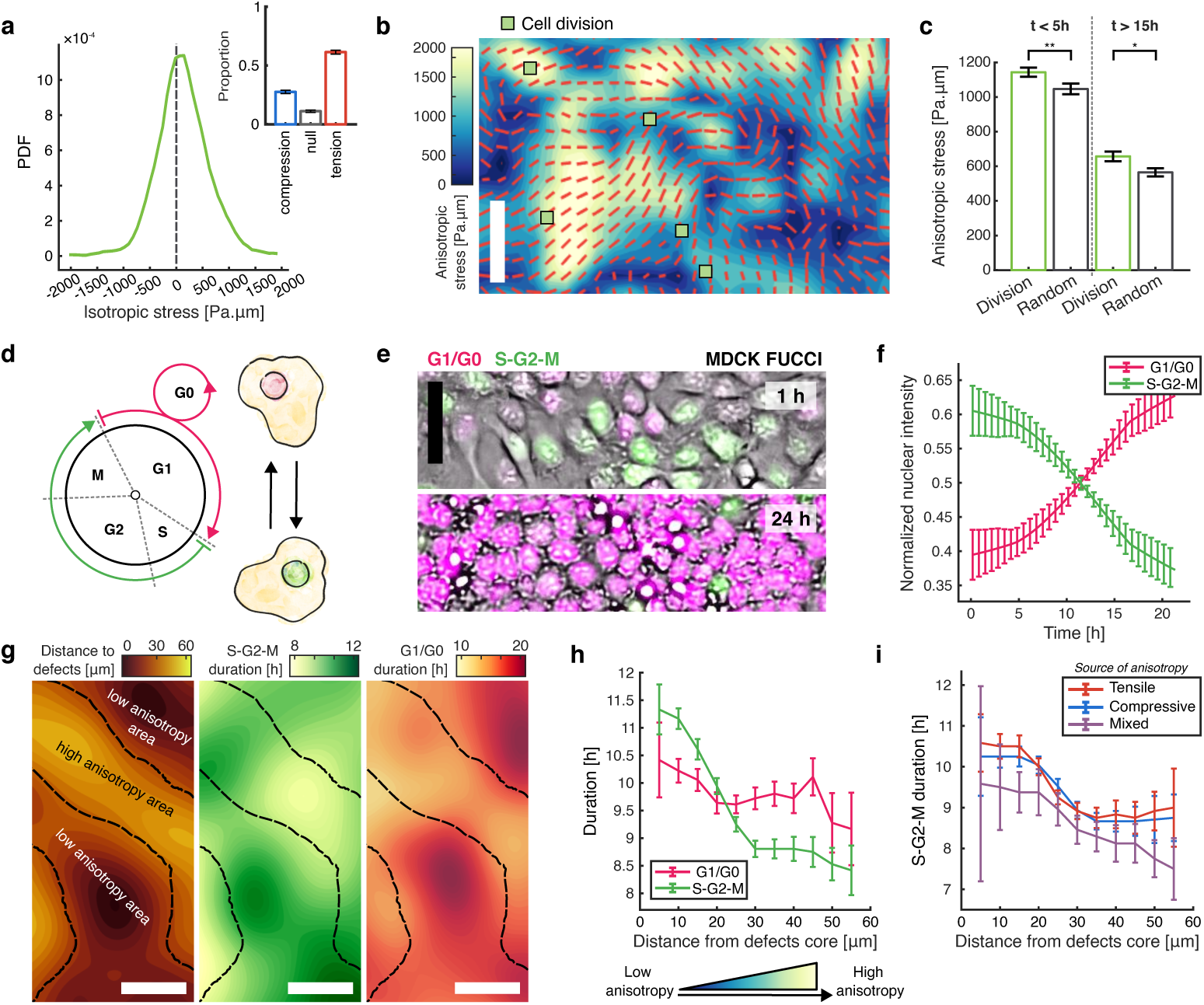
High anisotropic stress level modulates cell cycle completion. **a**) Probability distribution function (PDF) of the isotropic stress around cell division right before cytokinesis. 1672 cell divisions. Corresponding discrete barplot of the proportion of each mechanical state (compression, *σ_iso_ < −*100, tension, *σ_iso_ >* 100, and null, *−*100 *< σ_iso_ <* 100) is also shown. n = 13 movies from N = 3 independent experiments. **b**) Anisotropic stress field snapshot within the monolayer (t = 5h). Principal stress orientation in red. Cell divisions happening within a 1h window are marked in green square. **c**) Average anisotropic stress around cell division localization at the moment of cytokinesis and random position during densification. 583 divisions (t *<* 5h), 142 divisions (t *>* 15h), 266 random positions (t *<* 5h) and 387 random positions (t *>* 15h). n = 13 movies from N = 3 independent experiments. p *≤* 0.005 (t *<* 5h) and p *≤* 0.01 (t *>* 15h). **d**) Simplified diagram of the FUCCI MDCK cell line operating principle. The nuclear signal is pink when the cell is in G1 or G2 phase, and green when being in S, G2 or M phase. **e**) Top, phase contrast image of low density MDCK FUCCI monolayer (t = 1h) at low cell density, bottom, same monolayer at high cell density (t = 24h). Green is S-G2-M signalling, pink is G1/G0 signalling. **f**) Average evolution in time of the S-G2-M cells to total cell ratio (in green) and G1/G0 cells to total cell ratio (in pink) during tissue densification. N = 10 movies from n = 2 independent experiments. **g**) Left to right in order, temporal average of the distance from defects core map, corresponding map of the S-G2-M duration, projected in time, corresponding map of the G1/G0 duration, projected in time, and schematic of the corresponding expected anisotropic stress relative levels. The dashed black line corresponds to the isoline of a distance to defects of 31 µm. **h**) Cell cycle phase duration (green, S-G2-M, pink, G1/G0) regarding distance from stress defects core. N = 14 movies from n = 3 independent experiments. **i**) Cell cycle S-G2-M phase duration depending on the local anisotropic stress generation (blue, compressive anisotropy, red tensile anisotropy, mix of tensile and compressive anisotropy) regarding distance from stress defects core. N = 14 movies from n = 3 independent experiments. [Scale bar 50 µm (b, e, g). P-value from Mann-Whitney U-test (c). All error bars show the confidence interval at 95 %.]

Using MDCK FUCCI cells, we tracked cell cycle phases during tissue densification. As expected, increasing density correlated with more cells in G1/G0 phases, consistent with contact inhibition of proliferation (Fig. 3d–f, Movie S5). Yet local stress measurements revealed a striking pattern: cells in regions of high anisotropy exhibited shortened S–G2–M phases, resulting in faster cell cycle progression (Fig. 3g–h). This effect was observed regardless of the nature of the stress anisotropy, whether tensile, compressive, or a mixture of both. (Fig. 3i and Fig. 2a-b).

These results revise our understanding of mechanical regulation of proliferation. Rather than acting as a simple on–off switch based on scalar pressure, cells appear to respond to the degree of directional imbalance. Moreover, the results show that anisotropic stress can regulate the cell cycle regardless of the state of its isotropic counterpart. Importantly, these results do not contradict previous work, since isotropic stress can be associated with an anisotropic stress organization [8, 33]. This reframing resolves contradictions in the literature and positions anisotropy as a primary mechanical regulator of cell division.

### Cell division releases stress anisotropy through a feedback loop

If anisotropic stress promotes proliferation, how does the tissue avoid never-ending growth? Cell contact inhibition of proliferation is a well established mechanism in epithelia [11, 45], and increased cell density is often associated with elevated compressive stress [10, 12, 44]. In our data however, we did not observe an increase of isotropic compressive stress with tissue densification (Fig. S9a-c). Thus, we hypothesized that divisions themselves act back on the stress anisotropy, closing a feedback loop [15, 46].

Time-resolved stress maps confirmed this: cytokinesis was consistently followed by a local drop in anisotropy, whether cells divided under tension or compression (Fig. 4a). Both maximum and minimum principal stresses relaxed in a coordinated way (Fig. S10a), showing that divisions actively dissipate anisotropy along the division axis rather than passively reflecting it. Notably, we observed that when cells divided under tension, the compressive stress generated during cytokinesis counteracted the anisotropic tensile stress (Fig. 4b-c, [+] distribution). Alternatively, under compression, while the initial load acted perpendicular to the cell axis, cytokinesis generated compression along the division axis, thereby also reducing the difference between principal stresses (Fig. S10b-d) and thus decreasing stress anisotropy, despite increasing the local compression (Fig. 4b-c, [-] distribution).

**Fig. 4.**
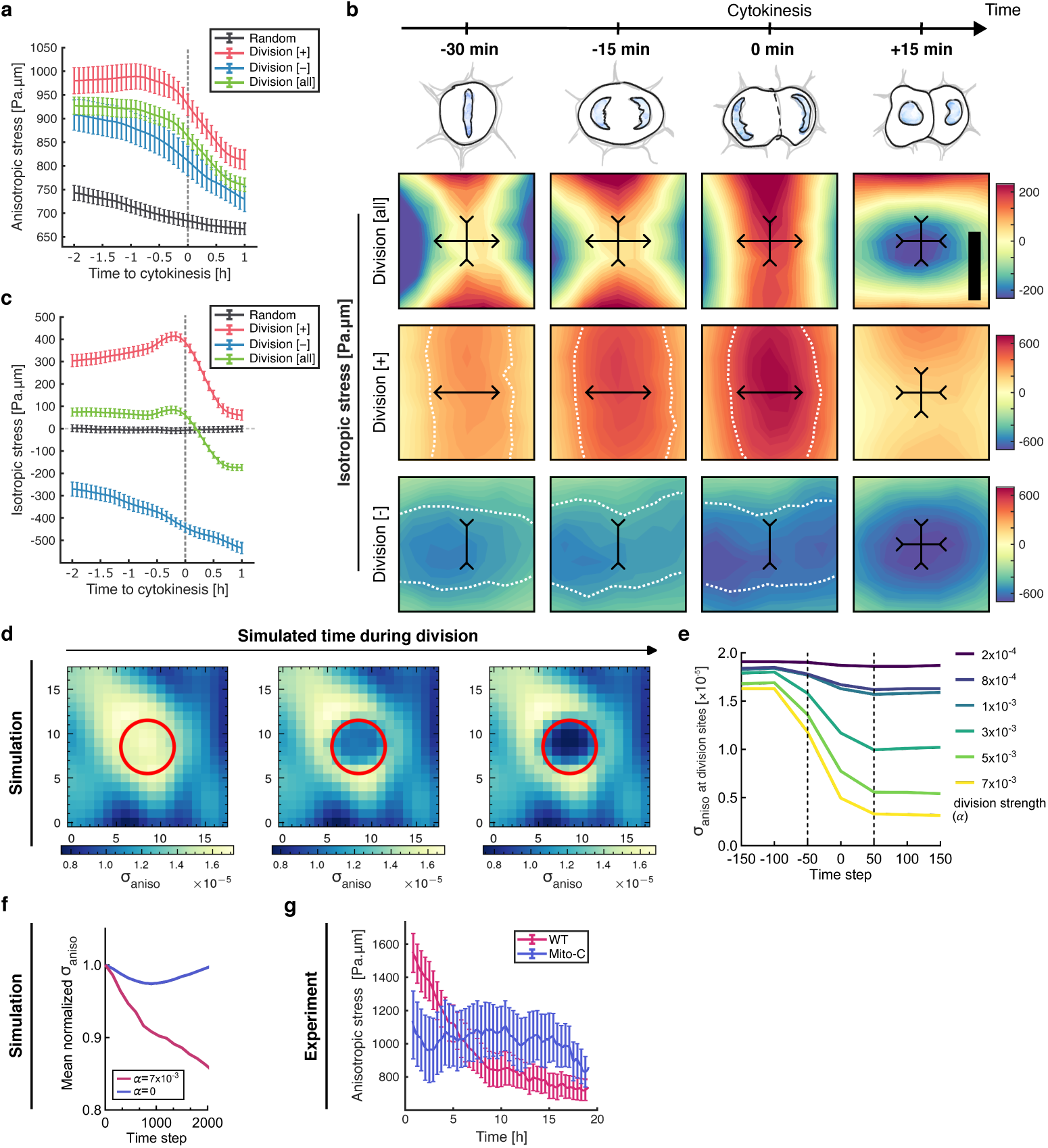
Cell division events release stress anisotropy during densification. **a**) Average anisotropic stress around cell undergoing mitosis. Cytokinesis happens at t = 0h. The legend show the average for random position, for division events undistinguished ([all]) and for sub-distribution of cells dividing under tension ([+]) and cell dividing under compression ([-]). 1672 cell divisions, 498 cell divisions [-], 826 cell divisions [+] and 2219 random positions). **b**) Isotropic stress field around cell division before and after cytokinesis, oriented along the cell division axis. The black double arrow show the orientation of the maximum and minimum stress, with their sign. top, 1478 cell divisions [all], middle 690 cell divisions [+], and bottom 457 cell divisions [-]. n = 13 movies from N = 3 independent experiments. **c**) Average isotropic stress around cell undergoing mitosis. Cytokinesis happens at t = 0h. The legend show the average for random position, for division events undistinguished ([all]) and for sub-distribution of cells dividing under tension ([+]) and cell dividing under compression ([-]). 1672 cell divisions [all], 498 cell divisions [-], 826 cell divisions [+] and 2219 random positions. n = 13 movies from N = 3 independent experiments. **d)** Simulated spatial maps of anisotropic stress before, during, and after a localized division event. The red circle indicates the site of division. **e)** Temporal evolution of anisotropic stress at the division site in simulations with varying division strengths (*α*). Black dashed lines denote the division period (Δ*t* = 100 time steps) **f)** Simulated domain-averaged normalized anisotropic stress over time for division-active (*α >* 0) and division-inhibited (*α* = 0) conditions. **g)** Experimental measurements of the average anisotropic stress within the whole tissue, for MDCK WT and for mitomycin-C treated MDCK cells. n = 24 movies from N = 5 independent experiments. [Scale bar 25 µm (b). All error bars show the confidence interval at 95 %.]

Simulations with a minimal active gel model [47, 48] captured this prediction: inserting artificial divisions at high-anisotropy sites triggered local stress relaxation (Fig. 4d–e and Fig. S11), while tissues without divisions remained mechanically frozen (Fig. 4f).

To test whether divisions are required for stress relaxation, we blocked proliferation using mitomycin-C. In these arrested monolayers, anisotropic stress no longer declined with time but instead accumulated and remained elevated (Fig. 4g). Moreover, by inhibiting proliferation at different stages of tissue densification, we were able to preserve distinct magnitudes of anisotropic stress corresponding to the level reached at the time of inhibition (Fig. S12a-c). This result rules out passive remodeling as the main driver of stress dissipation: without divisions, the feedback loop is broken and anisotropy persists. Thus, cytokinesis is not merely correlated with stress relaxation but is necessary for maintaining stress homeostasis.

To further investigate the role of cytokinesis, we perturbed cytokinetic contractility using increasing doses of blebbistatin (Fig. S12d-f). Consistent with our hypothesis, inhibition of contractile ring constriction impaired stress relaxation and led to the persistence of anisotropic stress, supporting a direct role for cytokinesis in the dissipation of mechanical anisotropy. Conversely, drug washout restored the feedback loop, enabling the monolayer to progressively relax accumulated stress as cell division resumed (Fig. S12g-i).

Together, these findings define a negative feedback loop: anisotropy promotes division, and division releases anisotropy (Fig. 5). This reciprocal coupling functions as a mechanical statos during growth: a feedback system in which stress anisotropy both drives and is dissipated by division, maintaining tissues in a dynamically stable state. Remarkably, this mechanism appears general and persists across different epithelial cell types (Fig. S13, Movie S6-7).

**Fig. 5.**
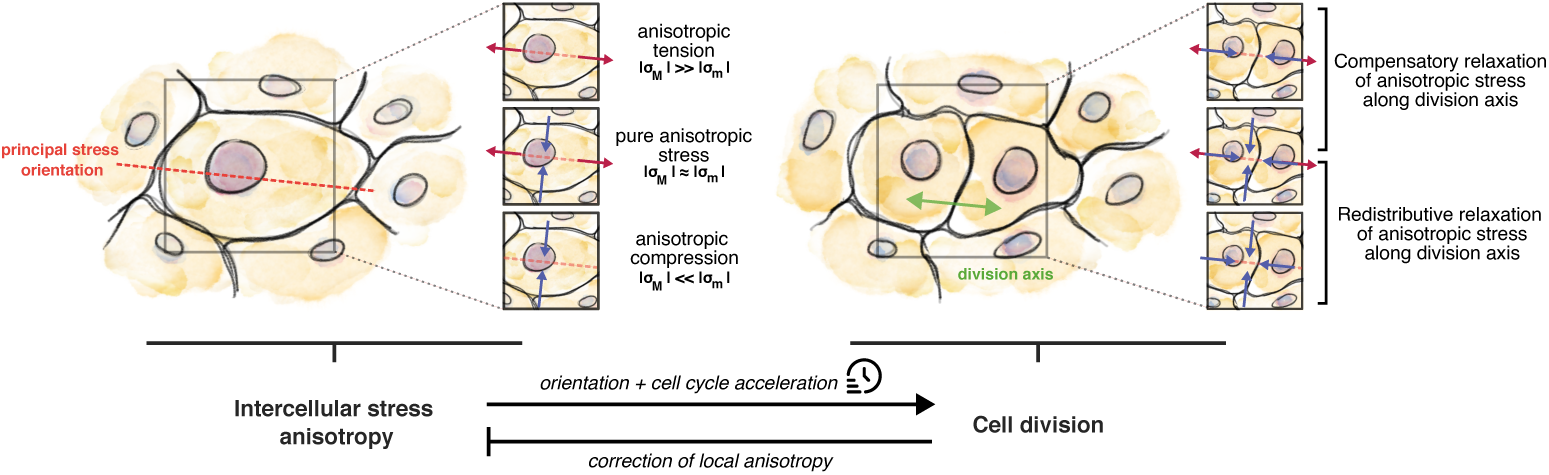
A principal-stress rule for cell division in epithelia. Cartoon summarizing the proposed mechanism : negative feed-back loop between anisotropic stress levels and cell proliferation in epithelia, across various coupling situation of principal stresses.

## Discussion

Taken together, these results identify a general physical principle governing how a dense active material relieves internal mechanical stress: through a discrete structural remodeling event, cell division, oriented along the axis of maximal stress anisotropy. When viewed alongside the literature (Supplementary Table 1), this principal-stress rule is not a special case but a genuine extension of the classical long-axis rule: in crowded tissues, where cell elongation and actin-based orientational order collapse, the orientation of structural remodeling is set not by geometry but by the direction of maximal stress anisotropy. This is consistent with previous reports that anisotropic tension can orient cell division [4, 13, 15, 28, 38, 49] even when it mismatches cell shape [14, 19, 20], and extends further to cases governed by anisotropic compression [6, 16, 17, 50, 51] or an unresolved isotropic stress state [3, 4]. That it remains predictive in vitro and ex vivo, across densities, cell types, and mechanical states, suggests this is a material property of dense epithelia, not a peculiarity of a single system.

Beyond orientation, anisotropy also regulates when cells divide by shortening S–G2–M, while cytokinesis dissipates anisotropy, forming a negative feedback that stabilizes the tissue’s mechanical state. Thus, division is both driven by and acts upon the stress landscape, providing a unified mechanical feedback loop for epithelial homeostasis (Fig. 5). Such coupling may represent a general strategy by which dense active materials avoid runaway internal stress while continuing to grow.

This loop provides a mechanical statos: proliferation is triggered by, and simultaneously relieves, stress anisotropy. Mitomycin-C and blebbistatin experiments confirmed the causal role of division in this balance, showing that when divisions were blocked or impaired, anisotropic stress remained elevated rather than relaxing. Such coupling may underlie how epithelia maintain growth while preventing mechanical overload.

Beyond this global homeostat, the stress field also contains finer-grained structures in the form of topological defects. Unlike shape- or actin-based defects, which vanish in crowded tissues, stress defects persist and correlate with division patterns: cells avoid dividing at defect cores but proliferate more frequently in surrounding regions of high anisotropy. These results highlight defects as potential organizing centers for division, opening the door to future studies of how stress defects interact with flows, tissue remodeling, and possibly even the defect dynamics familiar from active matter physics.

This principle has implications that extend beyond epithelial biology to the general mechanics of dense active and disordered materials. During development, anisotropic stress provides a robust orientational signal when geometric cues fluctuate, driving oriented remodeling that shapes tissue architecture [28, 43, 52]. In wound healing, stress anisotropy concentrated at wound edges [33, 53] may explain reoriented divisions that promote closure. In cancer, failure to relieve accumulated anisotropy may permit unchecked structural remodeling and invasive growth [45, 54]. And in engineered tissues, organoids, and synthetic active materials, deliberately imposed stress anisotropy could provide a route to program the geometry of structural remodeling, offering a design principle for materials that self-organize under internal mechanical control [42]. By shifting the organizing variable from geometry to stress, this work establishes a constitutive-like, testable rule for how dense active materials restructure under internal mechanical load, linking stress, structural remodeling, and homeostasis within a single quantitative framework.

## Acknowledgements.

We thank all the members of the “Cell Adhesion and Mechanics” team for helpful discussion, especially Yuan Shen, Joseph d’Alessandro and Gregory Arkowitz. We thank Tien Dang for establishing various MDCK cell lines, Joud Rachidi for helping with blebbistatin experiments, and Manon Arnaud for helping with the organoids culture. We thank James G. Rheinwald for allowing the use and Carien Niessen for sharing the N/TERT1 cell line. We acknowledge the ImagoSeine core facility of Institute Jacques Monod, member of France-BioImaging (ANR-10-INBS-04) and IBiSA, with support of Labex “Who Am I”, Inserm Plan Cancer, Region Ile-de-France and Fondation Bettencourt-Schueller.

## Funding

This work was supported by the European Research Council (Adv-101019835 “DeadorAlive” to B.L.), the Alexander von Humboldt Foundation (Alexander von Humboldt Professorship to B.L.), the Agence Nationale de la Recherche (“STRATEPI” DFG-ANR-22-CE92-0048 to R.M.M.), the Novo Nordisk Foundation (grant No. NNF18SA0035142 and NERD grant No. NNF21OC0068687 to A.D.), Villum Fonden (Grant no. 29476 to A.D.), and the European Union (ERC, PhysCoMeT, 101041418 to A.D.), the CNRS through 80 Prime program (to A.S. and B.L.) and Institut National du Cancer (INCa 18429 to R.M.M. and B.L.). L.A. received funding from the Ligue contre le Cancer and the Fondation pour la Recherche contre le Cancer. F.W. acknowledges financial support from the LabEx “Who Am I?” (grant ANR-11-LABX-0071). A.S. received funding from the Fondation Recherche Medicale (FDT-202404018282). Views and opinions expressed are those of the authors only and do not necessarily reflect those of the European Union or the European Research Council. Neither the European Union nor the granting authority can be held responsible for them.

## Author contributions

L.A., R.M.M., A.D. and B.L. designed the research. L.A. performed MDCK experiments and analyzed and interpreted all the experimental data. T.M. and K.T. established the model and performed simulations. F.W. performed MDCK immunostaining and helped with microscopy. A.S. performed keratinocyte experiments. S.K.M. performed intestinal organoids experiments. C.R. performed MCF10A experiments. M.A.F. contributed to the formal analysis and interpretations of the results. L.A., T.M., R.M.M., A.D. and B.L. wrote the manuscript. All authors read the manuscript and provided input on it.

## Competing interests

The authors declare no competing interests.

## Data and materials availability

All data supporting the findings of this study are included within the Article and its Supplementary Information. Source data are provided with this paper. All code used to process the data and to perform the experimental analysis are available upon reasonable request.

## Supplementary Materials

- Materials and Methods
- Supporting Information Text
- Supplementary Table 1
- References
- Supplementary Figures Fig. S1 to S13
- Movie S1 to S7

## Notes

### Competing Interest Statement

The authors have declared no competing interest.

## References

[1] Hertwig, O.: Welchen einfluß übt die schwerkraft auf die theilung der zellen? (2) (1884)

[2] Hertwig, O.: Ueber den werth der ersten furchungszellen für die organbildung des embryo experimentelle studien am frosch-und tritonei. Archiv für mikroskopische Anatomie 42(4), 662–807 (1893)

[3] Bosveld, F., Markova, O., Guirao, B., Martin, C., Wang, Z., Pierre, A., Balakireva, M., Gaugue, I., Ainslie, A., Christophorou, N., et al.: Epithelial tricellular junctions act as interphase cell shape sensors to orient mitosis. Nature 530(7591), 495–498 (2016)

[4] Nestor-Bergmann, A., Stooke-Vaughan, G.A., Goddard, G.K., Starborg, T., Jensen, O.E., Woolner, S.: Decoupling the Roles of Cell Shape and Mechanical Stress in Orienting and Cueing Epithelial Mitosis. Cell Reports 26(8), 2088–21004 (2019)

[5] Leen, E.V., Di Pietro, F., Bellaïche, Y.: Oriented cell divisions in epithelia: from force generation to force anisotropy by tension, shape and vertices. Current Opinion in Cell Biology 62, 9–16 (2020)

[6] Minc, N., Burgess, D., Chang, F.: Influence of cell geometry on division-plane positioning. Cell 144(3), 414–426 (2011)

[7] Gudipaty, S.A., Lindblom, J., Loftus, P.D., Redd, M.J., Edes, K., Davey, C., Krishnegowda, V., Rosenblatt, J.: Mechanical stretch triggers rapid epithelial cell division through piezo1. Nature 543(7643), 118–121 (2017)

[8] Uroz, M., Wistorf, S., Serra-Picamal, X., Conte, V., Sales-Pardo, M., Roca-Cusachs, P., Guimerà, R., Trepat, X.: Regulation of cell cycle progression by cell–cell and cell–matrix forces. Nature cell biology 20(6), 646–654 (2018)

[9] Curtis, A., Seehar, G.: The control of cell division by tension or diffusion. Nature 274(5666), 52–53 (1978)

[10] Di Meglio, I., Trushko, A., Guillamat, P., Blanch-Mercader, C., Abuhattum, S., Roux, A.: Pressure and curvature control of the cell cycle in epithelia growing under spherical confinement. Cell reports 40(8) (2022)

[11] Puliafito, A., Hufnagel, L., Neveu, P., Streichan, S., Sigal, A., Fygenson, D.K., Shraiman, B.I.: Collective and single cell behavior in epithelial contact inhibition. Proceedings of the National Academy of Sciences 109(3), 739–744 (2012)

[12] Devany, J., Falk, M.J., Holt, L.J., Murugan, A., Gardel, M.L.: Epithelial tissue confinement inhibits cell growth and leads to volume-reducing divisions. Developmental cell 58(16), 1462–1476 (2023)

[13] Fink, J., Carpi, N., Betz, T., Bétard, A., Chebah, M., Azioune, A., Bornens, M., Sykes, C., Fetler, L., Cuvelier, D., et al.: External forces control mitotic spindle positioning. Nature cell biology 13(7), 771–778 (2011)

[14] Scarpa, E., Finet, C., Blanchard, G.B., Sanson, B.: Actomyosin-driven tension at compartmental boundaries orients cell division independently of cell geometry in vivo. Developmental cell 47(6), 727–740 (2018)

[15] Wyatt, T.P., Harris, A.R., Lam, M., Cheng, Q., Bellis, J., Dimitracopoulos, A., Kabla, A.J., Charras, G.T., Baum, B.: Emergence of homeostatic epithelial packing and stress dissipation through divisions oriented along the long cell axis. Proceedings of the National Academy of Sciences 112(18), 5726–5731 (2015)

[16] Blanchard, G.B., Scarpa, E., Muresan, L., Sanson, B.: Mechanical stress combines with planar polarised patterning during metaphase to orient embryonic epithelial cell divisions. Development 151(10), 202862 (2024)

[17] Middelkoop, T.C., Neipel, J., Cornell, C.E., Naumann, R., Pimpale, L.G., Jülicher, F., Grill, S.W.: A cytokinetic ring-driven cell rotation achieves hertwig’s rule in early development. Proceedings of the National Academy of Sciences 121(25), 2318838121 (2024)

[18] Shroff, N.P., Xu, P., Kim, S., Shelton, E.R., Gross, B.J., Liu, Y., Gomez, C.O., Ye, Q., Drennon, T.Y., Hu, J.K., et al.: Proliferation-driven mechanical compression induces signalling centre formation during mammalian organ development. Nature cell biology 26(4), 519–529 (2024)

[19] Finegan, T.M., Na, D., Cammarota, C., Skeeters, A.V., Nádasi, T.J., Dawney, N.S., Fletcher, A.G., Oakes, P.W., Bergstralh, D.T.: Tissue tension and not interphase cell shape determines cell division orientation in the drosophila follicular epithelium. The EMBO journal 38(3), 100072 (2019)

[20] Hart, K.C., Tan, J., Siemers, K.A., Sim, J.Y., Pruitt, B.L., Nelson, W.J., Gloerich, M.: E-cadherin and lgn align epithelial cell divisions with tissue tension independently of cell shape. Proceedings of the National Academy of Sciences 114(29), 5845–5853 (2017)

[21] Bi, D., Yang, X., Marchetti, M.C., Manning, M.L.: Motility-driven glass and jamming transitions in biological tissues. Physical Review X 6(2), 021011 (2016)

[22] Devany, J., Sussman, D.M., Yamamoto, T., Manning, M.L., Gardel, M.L.: Cell cycle–dependent active stress drives epithelia remodeling. Proceedings of the National Academy of Sciences 118(10), 1917853118 (2021)

[23] Atia, L., Bi, D., Sharma, Y., Mitchel, J.A., Gweon, B., A. Koehler, S., DeCamp, S.J., Lan, B., Kim, J.H., Hirsch, R., et al.: Geometric constraints during epithelial jamming. Nature physics 14(6), 613–620 (2018)

[24] Stooke-Vaughan, G.A., Campàs, O.: Physical control of tissue morphogenesis across scales. Current opinion in genetics & development 51, 111–119 (2018)

[25] Mao, Y., Wickström, S.A.: Mechanical state transitions in the regulation of tissue form and function. Nature Reviews Molecular Cell Biology 25(8), 654–670 (2024)

[26] Saw, T.B., Doostmohammadi, A., Nier, V., Kocgozlu, L., Thampi, S., Toyama, Y., Marcq, P., Lim, C.T., Yeomans, J.M., Ladoux, B.: Topological defects in epithelia govern cell death and extrusion. Nature 544(7649), 212–216 (2017)

[27] Balasubramaniam, L., Monfared, S., Ardaševa, A., Rosse, C., Schoenit, A., Dang, T., Maric, C., Hautefeuille, M., Kocgozlu, L., Chilupuri, R., et al.: Dynamic forces shape the survival fate of eliminated cells. Nature physics 21(2), 269–278 (2025)

[28] Campinho, P., Behrndt, M., Ranft, J., Risler, T., Minc, N., Heisenberg, C.-P.: Tension-oriented cell divisions limit anisotropic tissue tension in epithelial spreading during zebrafish epiboly. Nature cell biology 15(12), 1405–1414 (2013)

[29] Nelson, C.M., Xiao, B., Wickström, S.A., Dufrêne, Y.F., Cosgrove, D.J., Heisenberg, C.-P., Dupont, S., Shyer, A.E., Rodrigues, A.R., Trepat, X., et al.: Mechanobiology: Shaping the future of cellular form and function. Cell 187(11), 2652–2656 (2024)

[30] Ladoux, B., Mège, R.-M.: Mechanobiology of collective cell behaviours. Nature reviews Molecular cell biology 18(12), 743–757 (2017)

[31] Nier, V., Jain, S., Lim, C.T., Ishihara, S., Ladoux, B., Marcq, P.: Inference of internal stress in a cell monolayer. Biophysical journal 110(7), 1625–1635 (2016)

[32] Anger, L., Schoenit, A., Wodrascka, F., Rossé, C., Mège, R.-M., Ladoux, B., Marcq, P.: Tissue stress measurements with bayesian inversion stress microscopy. The European Physical Journal E 49(1), 2 (2026)

[33] Tambe, D.T., Corey Hardin, C., Angelini, T.E., Rajendran, K., Park, C.Y., Serra-Picamal, X., Zhou, E.H., Zaman, M.H., Butler, J.P., Weitz, D.A., et al.: Collective cell guidance by cooperative intercellular forces. Nature materials 10(6), 469–475 (2011)

[34] Sato, T., Vries, R.G., Snippert, H.J., Van De Wetering, M., Barker, N., Stange, D.E., Van Es, J.H., Abo, A., Kujala, P., Peters, P.J., et al.: Single lgr5 stem cells build crypt-villus structures in vitro without a mesenchymal niche. Nature 459(7244), 262–265 (2009)

[35] Pérez-González, C., Ceada, G., Greco, F., Matejčić, M., Gómez-González, M., Castro, N., Menendez, A., Kale, S., Krndija, D., Clark, A.G., et al.: Mechanical compartmentalization of the intestinal organoid enables crypt folding and collective cell migration. Nature cell biology 23(7), 745–757 (2021)

[36] Malinverno, C., Corallino, S., Giavazzi, F., Bergert, M., Li, Q., Leoni, M., Disanza, A., Frittoli, E., Oldani, A., Martini, E., et al.: Endocytic reawakening of motility in jammed epithelia. Nature materials 16(5), 587–596 (2017)

[37] Shen, Y., O’Byrne, J., Schoenit, A., Maitra, A., Mège, R.-M., Voituriez, R., Ladoux, B.: Flocking and giant fluctuations in epithelial active solids. Proceedings of the National Academy of Sciences 122(16), 2421327122 (2025)

[38] LeGoff, L., Rouault, H., Lecuit, T.: A global pattern of mechanical stress polarizes cell divisions and cell shape in the growing drosophila wing disc. Development 140(19), 4051–4059 (2013)

[39] Duclos, G., Erlenkämper, C., Joanny, J.-F., Silberzan, P.: Topological defects in confined populations of spindle-shaped cells. Nature Physics 13(1), 58–62 (2017)

[40] Kawaguchi, K., Kageyama, R., Sano, M.: Topological defects control collective dynamics in neural progenitor cell cultures. Nature 545(7654), 327–331 (2017)

[41] Park, J.-A., Kim, J.H., Bi, D., Mitchel, J.A., Qazvini, N.T., Tantisira, K., Park, C.Y., McGill, M., Kim, S.-H., Gweon, B., et al.: Unjamming and cell shape in the asthmatic airway epithelium. Nature materials 14(10), 1040–1048 (2015)

[42] Guillamat, P., Mirza, W., Bal, P.K., Gómez-González, M., Roca-Cusachs, P., Arroyo, M., Trepat, X.: Guidance of cellular nematic elastomers into shape-programmable living surfaces. Science 392(6795), 317–323 (2026)

[43] Cislo, D.J., Yang, F., Qin, H., Pavlopoulos, A., Bowick, M.J., Streichan, S.J.: Active cell divisions generate fourfold orientationally ordered phase in living tissue. Nature physics 19(8), 1201–1210 (2023)

[44] Donker, L., Houtekamer, R., Vliem, M., Sipieter, F., Canever, H., Gomez-Gonzalez, M., Bosch-Padros, M., Pannekoek, W.-J., Trepat, X., Borghi, N., et al.: A mechanical g2 checkpoint controls epithelial cell division through e-cadherin-mediated regulation of wee1-cdk1. Cell reports 41(2) (2022)

[45] McClatchey, A.I., Yap, A.S.: Contact inhibition (of proliferation) redux. Current opinion in cell biology 24(5), 685–694 (2012)

[46] Gupta, V.K., Nam, S., Yim, D., Camuglia, J., Martin, J.L., Sanders, E.N., O’Brien, L.E., Martin, A.C., Kim, T., Chaudhuri, O.: The nature of cell division forces in epithelial monolayers. Journal of Cell Biology 220(8), 202011106 (2021)

[47] Doostmohammadi, A., Thampi, S.P., Yeomans, J.M.: Defect-mediated morphologies in growing cell colonies. Physical review letters 117(4), 048102 (2016)

[48] Thampi, S.P., Doostmohammadi, A., Golestanian, R., Yeomans, J.M.: Intrinsic free energy in active nematics. Europhysics Letters 112(2), 28004 (2015)

[49] Louveaux, M., Julien, J.-D., Mirabet, V., Boudaoud, A., Hamant, O.: Cell division plane orientation based on tensile stress in arabidopsis thaliana. Proceedings of the National Academy of Sciences 113(30), 4294–4303 (2016)

[50] Lisica, A., Fouchard, J., Kelkar, M., Wyatt, T.P., Duque, J., Ndiaye, A.-B., Bonfanti, A., Baum, B., Kabla, A.J., Charras, G.T.: Tension at intercellular junctions is necessary for accurate orientation of cell division in the epithelium plane. Proceedings of the National Academy of Sciences 119(49), 2201600119 (2022)

[51] Lintilhac, P.M., Vesecky, T.B.: Stress-induced alignment of division plane in plant tissues grown in vitro. Nature 307(5949), 363–364 (1984)

[52] Streichan, S.J., Hoerner, C.R., Schneidt, T., Holzer, D., Hufnagel, L.: Spatial constraints control cell proliferation in tissues. Proceedings of the National Academy of Sciences 111(15), 5586–5591 (2014)

[53] Chen, T., Callan-Jones, A., Fedorov, E., Ravasio, A., Brugués, A., Ong, H.T., Toyama, Y., Low, B.C., Trepat, X., Shemesh, T., et al.: Large-scale curvature sensing by directional actin flow drives cellular migration mode switching. Nature physics 15(4), 393–402 (2019)

[54] Matthews, H.K., Bertoli, C., Bruin, R.A.: Cell cycle control in cancer. Nature reviews Molecular cell biology 23(1), 74–88 (2022)

