## Supplementary Information for "A principal-stress rule for cell division in epithelia"

**The PDF file includes:**

- Materials and Methods
- Supporting Information Text
- Supplementary Table 1
- References
- Supplementary Figures Fig. S1 to S13

**Other Supplementary Materials for this manuscript include the following:**

- Movie S1 to S7

### Materials and Methods

#### Cell lines culture

The following cell lines were used:

- MDCK-II (ATCC CCL-34)
- MDCK-II CAAX GFP (recently described in [1])
- MDCK-II FUCCI cell cycle reporter (described in [2])
- MCF10A (ATCC CRL-10317)
- N/TERT1 keratinocytes (gift from the Niessen laboratory (CECAD Cologne, Germany, generated by the Rheinwald laboratory, Harvard), first described in [3]).

MDCK-II WT, MDCK-II FUCCI and MDCK-II CAAX GFP cells were cultured in DMEM (GlutaMAX, supplemented with high glucose and pyruvate, Life Technologies), with 10% fetal bovine serum (FBS, Life Technologies) and 1% penicillin-streptomycin (Life Technologies) at 37°C with 5% $CO_2$. MCF-10A cells were grown in DMEM-F12 (Gibco) containing 10% penicillin-glutamine, 10 $\mu g/mL$ human insulin (Sigma-Aldrich), 100 ng/mL cholera toxin (Sigma-Aldrich), 0.5 mg/mL hydrocortisone (Sigma-Aldrich), 5% horse serum, and 20 ng/ml EGF (PeproTeck) at 37°C with 5% $CO_2$. Keratinocytes were maintained in growth medium (CnT-Prime, CellnTech, Switzerland) at 37°C with 5% $CO_2$, and switched to differentiation medium (supplemented with high calcium and growth factors, CnT-Prime-3D, CellnTech, Switzerland) 2 days before imaging. Cell populations were split using 0. 05% trypsin (Merck) twice a week. Before processing, the culture medium was aspirated and cells were rinsed with PBS to remove dead cells and debris. Mycoplasma contamination in cell cultures was routinely checked using a PCR test.

#### Organoid preparation and culture

Wild-type C57/Bl6 6–12 weeks-old male mice were provided by the animal house facility of the Institut Jacques Monod, and were used for organoid generation. Following euthanasia by cervical dislocation, the small intestine was collected, rinsed with PBS to remove luminal contents, and opened longitudinally. The tissue was then cut into 3–5 mm fragments and washed again in PBS. Organoid preparation and culture was done exactly following the method described in [4, 5].

H2B-mCherry organoids were generated from mice provided by Renata Basto (Institut Curie, Paris)[6]. VillinCreERT2-tdTomato organoids were generated from mice provided by Danijela Vignjevic (Insitut Curie, Paris)[7].

#### Sample preparation

For TFM experiments, cells were grown for 15–20 hours until they formed a monolayer at 90–100% confluency. Prior to imaging, the culture medium was aspirated, and cells were rinsed with PBS to remove dead cells and debris. For TFM experiments involving inhibition of cell division, cells that reached 95% confluency were incubated with mitomycin-C (Sigma-Aldrich) at a concentration of 20 µM for 45 minutes. The medium was then removed, cells were rinsed with PBS, and incubation continued for 12–15 hours to allow all cells to reach the mitotic arrest stage induced by mitomycin-C, prior to imaging. For experiments at different stade of densification, the same protocol was applied, but mitomycin was simply added X days after reaching 95% confluency (X being equal to either 1, 2 or 3). For culture of organoids in 2D, 3D organoids were cultured in conditioned media for at least 3 days before use, mechanically broken and used to generated 2D monolayers following the method used in [5, 8]. The organoid-derived monolayers were cultured for 10 days prior live imaging. For TFM experiments involving inhibition of myosin-II, cells that reached 100% confluency were incubated with blebbistatin (Sigma-Aldrich), at either 10, 20, or 50 $\mu$M. Control were realized with a corresponding volume of 50 $\mu$M DMSO.

For fixed staining, cells that reached the desired density were first rinsed three times with PBS, then incubated with 4% paraformaldehyde (PFA) for 20 minutes at room temperature. Cells were subsequently permeabilized with 0.5% Triton X-100 for 20 minutes. After three 5-minute washes in PBS, samples were blocked with 1% bovine serum albumin (BSA) and 10% fetal bovine serum (FBS) in PBS for 1 hour at room temperature. Samples were then washed three times in PBS, and the actin cytoskeleton was stained using either phalloidin–Alexa Fluor 488 (Invitrogen) or phalloidin–Alexa Fluor 568 (Life Technologies), diluted at 0.5% in PBS. Nuclei were stained with Hoechst (Thermo Fisher), diluted at 0.05% in PBS, for 3 hours at room temperature. Samples were stored at 4°C until imaging.

#### Traction force protocol

Soft biocompatible polymer coated substrates for TFM were prepared as previously described [9]. Briefly, CY52-276A and CY52-276B polydimethylsiloxane (PDMS, Dowsil, Dow) were mixed using different weight ratios (1:1 to achieve 15 kPa, 5:6 to achieve 30kPa) and then poured into microfabrication polystyrene petri dishes (Bio-One, Greiner). The prepared surfaces were spin-coated for 30 seconds at 500 rpm in order to obtain a layer of thickness around 100 µm and then cured at 80°C for 2 hours. Once cured, each dish was silanized using a 10% APTES ((3-aminopropyl)triethoxysilane, Sigma-Aldrich) solution diluted in absolute ethanol for 15 min and then washed with absolute ethanol 3 times before being dried at 80°C for 10 min. 200 nm Cy3 red carboxylated fluorescent beads (FluoSpheres, Invitrogen) were diluted in water with a ratio of 2:1000 and disaggregated using an ultrasonic bath for 10 min. The beads solution was filtered using a 0.45 µm filter and incubated on TFM substrates for 15 min, protected from light. The dishes were washed with water 3 times and dried at 80°C for 3 minutes, and proper homogeneous deposition was checked under microscope. Prior to cell seeding, prepared substrates were coated with 50 µg/mL fibronectin (Sigma) for 45 min and washed 3 times with PBS. Cells were imaged for 24h and beads resting positions were obtained at the end of each experiment by adding 200 µL of 10% sodium dodecyl sulfate (SDS) in the medium in order to remove the cells.

For organoids experiments, 5kPa soft polyacrylamide (PAA) were used and prepared according to [4]. Briefly, glass coverslips were cleaned in an ethanol bath, sonicated for 1 min, dried at 80 °C for 15 min, and then exposed to plasma cleaning for 10 min. Afterward, the coverslips were incubated for 30 min in a silane solution containing 2% 3-(trimethoxysilyl)propylmethacrylate (Sigma-Aldrich, no. 440159) and 1% acetic acid in ethanol. The silanized coverslips were rinsed with ethanol, dried at 80°C for 1 h, and stored at room temperature. A freshly prepared polyacrylamide gel mixture (acrylamide, bis-acrylamide, ammonium persulphate, and TEMED in PBS, proportions as described in [4]) containing 4% fluorescent beads (FluoSpheres, Invitrogen) was cast onto the coverslips and allowed to polymerize at room temperature for 1h to form a 100-µm-thick gel. The resulting substrates were stored in PBS at 4 °C. Before use, PAA gels were functionalized with Sulpho-SANPAH (Cultek), activated under 365-nm UV light for 10 min, and coated with a solution of rat-tail type I collagen (First Link UK) and laminin-1 (Sigma-Aldrich) in PBS, followed by overnight incubation at 4 °C, as described in [8].

#### Microscopy and data visualization

Time-lapse traction force microscopy (TFM) was performed using a live-cell epifluorescence microscope (BioStation IM-Q, Nikon or Nikon confocal CSU), equipped with either a 10× or 20× phase-contrast air objective and a standard incubation chamber. Cells were maintained at 37°C with 5% $CO_2$. The monolayer was imaged using either phase contrast or DIC, and fluorescent beads were visualized using epifluorescence. For cell shape characterization, MDCK CAAX-GFP cells were used and imaged using epifluorescence. Time-lapse videos were recorded at multiple positions every 5 minutes (MDCK) or every 10 minutes (MCF10A and keratinocytes), for a typical duration of approximately 24 hours.

Confocal microscopy was performed using a laser scanning confocal microscope (ZEISS LSM 980), equipped with a 40× or 63× oil-immersion objective and an Airyscan 2 module. MDCK WT cells were grown, fixed and mounted on a glass-bottom imaging dish upon reaching the desired cell density. Unless otherwise specified, all images or Z-stacks were acquired in Airyscan mode without signal averaging, and automated deconvolution was performed using the microscope software (ZEISS ZEN Blue).

Most images were visualized using ImageJ [10], with brightness and contrast adjusted as needed. For Z-stacks, maximum intensity projections were generated. In cases where physical parameters such as the nematic director or principal stress orientation were overlaid, images were visualized using MATLAB. Brightness and contrast adjustments were also performed in MATLAB using the Image Processing Toolbox.

#### Analysis method

Unless stated otherwise, all codes for the following experimental analysis were written in MATLAB using built-in functions, standard toolboxes, or previously published tools implemented in MATLAB.

##### Shape characterization

Geometrical characterization of the cells during monolayer packing was performed using from MDCK CAAX-GFP cells signal. Cells were segmented using Cellpose with the pretrained model `cyto3` [11]. Customized binary images of the segmented cells were generated (see Fig. S4A), and a filter was applied based on the average area of each object to discard falsely detected cells, using the `regionprops` function of MATLAB. Cells with an area smaller than 80 $\mu m^2$ were excluded. The area, perimeter, aspect ratio (via ellipse fitting), and center-of-mass position were computed for each remaining object, post filtering. Using the formula $\rho_0 = \frac{P}{\sqrt{A}}$, where $P$ is the perimeter and $A$ is the area of the cell, the cell shape index $\rho_0$ was also computed. The center-of-mass positions were used to calculate the average distance between neighboring cells using a Euclidean metric.

#### Collective cell dynamic

Dynamical characterization during monolayer densification was performed by tracking individual cell movements within the monolayer. Cell tracking was conducted using TrackMate [12] on brightfield time-lapse images, employing a LoG detector combined with a median filter for cell detection, followed by an advanced Kalman tracker for tracking. The initial search radius was set to 15 $\mu m$ and allowed to increase up to 20 $\mu m$. Tracks were permitted to have one undetected frame. Each track provided the time-dependent center-of-mass position $\boldsymbol{r}_i(t) = [x_i(t), y_i(t)]$ for a given object $i$, enabling computation of the mean square displacement (MSD) as:

$$MSD(\Delta t) = \left\langle \|\boldsymbol{r}_i(t + \Delta t) - \boldsymbol{r}_i(t)\|^2 \right\rangle$$

where the average $\langle \cdot \rangle$ is taken over all cells $i$ and all time points $t$. Tracks were grouped into five time windows of 5 hours each (0–5 h, 5–10 h, etc.), and the MSD was computed over each period. The global cellular velocity field was quantified using the Particule Image Velocimetry toolbox PIVlab [13]. Displacement field was computed using an interrogation window of 64×64 pixels with 50% overlap. The resulting vector field was first smoothed using a second interrogation window of 32×32 pixels, and then using the built-in averaging statistical algorithm in the plugin. Velocity field was then computed accordingly, knowing the duration between two consecutive frames.

#### Nematic analysis

The nematic director field was approximated using intensity gradients within each cell, excluding cell edges. To compute the intensity gradient, the OrientationJ plugin [14] in ImageJ was used. The resulting field was coarse-grained using a moving square window of 100 pixels with 64 pixels of overlap. For visualization, the field was interpolated so that each director approximately corresponds to one cell.

Nematic topological defects were detected using a winding number approach, following a previously established pipeline [15–17]. For each detected $+\frac{1}{2}$ or $-\frac{1}{2}$ topological defect, the local divergence of the nematic director field was used to determine the orientation of the defect, as proposed in [18].

To quantify the loss of nematic ordering as cells lose their elongated shape as cell density increased, we defined an "apparent" scalar order parameter $q_a$ (Fig. S5C) as $q_a = \langle \nu(\boldsymbol{r})\, q(\boldsymbol{r}) \rangle$ where $q(\boldsymbol{r})$ is the local classical nematic scalar order parameter at the position $\boldsymbol{r}$, computed using a local window $w$ such as:

$$q(\boldsymbol{r}) = 2\cos^2(\theta_{\boldsymbol{r}} - \theta_w) - 1$$

Here, $\theta_{\boldsymbol{r}}$ is the local director orientation at the position $\boldsymbol{r}$, and $\theta_w$ is the average director field orientation within the window $w$. In our experiments, $w$ is a square of size 50 $\mu$m. The function $\nu(\boldsymbol{r})$ acts as a pseudo-flag based weight using the local shape index $\rho_0(\boldsymbol{r})$, and is defined as:

$$\nu(\boldsymbol{r}) = \begin{cases} 1.5 & \text{if } \rho_0(\boldsymbol{r}) > 4 \\ 0 & \text{if } \rho_0(\boldsymbol{r}) < 3.81 \\ 1 & \text{otherwise} \end{cases}$$

where $\rho_0(\boldsymbol{r})$ is the local shape index of the cell at the position $\boldsymbol{r}$. Simply put, the scalar order parameter was weighted by the local shape index to reduce its contribution in regions where cells adopt a more hexagonal morphology, thereby capturing the loss of nematic order during tissue densification.

#### Traction forces and stress measurement

Images of bead displacements relative to their relaxed positions were preprocessed using the Image Stabilizer plugin in ImageJ [19], and background illumination was corrected to reduce noise. The beads displacement field was obtained using PIVlab [13], with an interrogation window of 32×32 pixels and a 50% overlap. The corresponding traction force field was then inferred using Fourier transform traction cytometry, knowing theoretical substrate stiffness (15 or 30 kPa in our experiments) and a regularization parameter of $9\times10^{-9}$. From the traction force field, we computed the 2D stress tensor across the tissue using BISM [20], with a regularization parameter of $\Lambda = 10^{-6}$. The 2D stress tensor was then diagonalized in order to compute the principal stress direction as well as the maximum and the minimum principal stress, $\sigma_M$ and $\sigma_m$ (see Supporting Information).

Because principal stress orientation can be mathematically computed even in purely isotropic regions where principal stress orientation lacks physical meaning, we defined an apparent scalar order parameter for the stress tensor, $q'_a$, similarly to the nematic case. Here, the weighting factor is defined as $\nu' = \sqrt{3}\frac{\sigma_{aniso}}{\sigma_{VM}}$ where $\sigma_{VM}$ is the von Mises stress such as $\sigma_{VM} = \sqrt{\sigma_M^2 + \sigma_m^2 - \sigma_M\sigma_m}$. Defined that way, this anisotropic ratio (Fig. S9d-e) is bounded between 0 (when $\sigma_M = \sigma_m$) and 1 (when $\sigma_M = -\sigma_m$). The apparent stress scalar order parameter $q'_a$ was then defined as $q'_a = \langle \nu'(\boldsymbol{r})\, q'(\boldsymbol{r})\rangle$ with $q'(\boldsymbol{r})$ the local scalar order parameter computed using the director of the principal stress directions.

To detect stress topological defects, we used the same pipeline described above for nematic topological defects. To track the movement of stress defects over time, we generated movies in which the positions of each defect were represented as white objects on a black background. These binary position images were then analyzed using TrackMate, and all parameters used to characterize the motion of each trajectory were retrieved from the TrackMate analysis. Defect lifetime was defined as the duration of the track. The net displacement was computed as $d_{net} = \|\boldsymbol{r_f} - \boldsymbol{r_0}\|$ where $\boldsymbol{r_f}$ and $\boldsymbol{r_0}$ are the final and initial positions of the considered defect. The confinement ratio was defined as $CR = \frac{d_{net}}{d_{tot}}$ with $d_{tot}$ being the total cumulated distance traveled by the considered defect, and $d_{net}$ the actual net distance traveled by the considered defect. A confinement ratio approaching zero indicates restricted (caged) displacement, whereas a ratio approaching one corresponds to free rectilinear displacement. A binary spatial map of all cumulative defect positions (independent of topological charge) was then generated for each video, and these maps were subsequently used to compute the average spatial distribution of distances from defect cores.

#### Micropatterning and laser ablation

Microfabrication was done as previously described [9, 16, 21]. Mold of ellipse dimensioned as 150 x 300 $\mu$m were obtained using standard lithography methods. PDMS (SYLGARD 184, Dow Corning) was prepared by mixing the base with a curing agent at a ratio of 1:10, poured over the mold, degassed and then cured at 80○C for 2 h. Stamps were peeled of the mold and stored, protected from light and humidity. When utilized, a fibronectin solution (50 $\mu$ g/mL, Sigma) was incubated covering the surface of the mold for 45 min. The surface was cleaned using a gentle air flow, and the stamps were gently pressed against the bottom of a PDMS substrate for about 1 min. Patterns were then incubated with a solution of 2% Pluronic F-127 (Sigma) for 1 h to passivate the areas outside the patterns.

MDCK CAAX GFP cells were seeded on PDMS patterned dish and grown to confluency inside each ellipse. Before wound induction, dishes were rinsed using warm PBS and provided with a fresh culture medium. Laser ablation was done using a spinning-disk CSU-X1 with a fluorescence recovery after photo bleaching module (Yokogawa) and a ×40/1.2 water immersion objective. Briefly, ellipse were ablated along their central short axis by focusing an ultraviolet laser (355 nm, pulse duration 3–5 ns, laser power 450 mW) for 1 s. Each sample was imaged during 5 s before ablation and until 1 min after ablation, using 1s intervals. The recoil velocity was measured by manually segmenting the wound separation in the direction of the ellipse long axis over time and plotting the corresponding size rate evolution.

#### Cell division event

For MDCK WT, N/TERT1 keratinocytes, MCF10A and organoids data, position and orientation of cytokinesis events were manually detected in a homogeneous manner, without prior knowledge of the mechanical state of the tissue. To ensure a spatially uniform distribution of marked events, each image was divided into four quadrants, and an equal number of division events were marked in each. The typical number of divisions annotated per position in a given experiment was 125 for MDCK, 80 for the model systems. When a parameter was analyzed around a cell division event, it was generally evaluated locally at the position of the event. In Fig. 4 and Supplementary Fig. S10, however, the stresses were computed as the median within a 50 $\mu m$ square window centered on the event to provide a quantitative measure representative of the local stress patterns displayed in these figures.

For MDCK FUCCI data, binary masks for each signal (G1/G0 and S/G2/M) were generated after preprocessing using MATLAB's Image Processing Toolbox. Specifically, the `imbinarize` function was applied, followed by morphological operations such as erosion and dilation. Objects too small to represent nuclei were filtered based on area thresholds. Only complete trajectories, corresponding to cells entering and exiting a given FUCCI state within the imaging interval, were retained for analysis. Each binary image was analyzed using TrackMate to determine the lifetime of each detected object, corresponding by construction to the duration of a particular cell cycle phase. A two-dimensional spatial map of each phase duration was then constructed by assigning to each position **r** the average duration of all tracked cells observed at that location throughout the experiment. Formally, for a given spatial map of a considered cell cycle phase duration $D^{\tau}_{phase}$, $D^{\tau}_{phase}$ is computed as $D^{\tau}_{phase}(\boldsymbol{r}) = \langle \tau_{phase}(\boldsymbol{r}) \rangle$ where $\langle \cdot \rangle$ denotes the average over cell cycle duration $\tau_{phase}$ of the considered cell cycle phase at the position $\boldsymbol{r}$ during the course of experiment. A similar approach was adopted to characterize the average isotropic mechanical state experienced by cells over the course of the experiment. The isotropic stress field was first averaged over time, yielding a two-dimensional map of the mean isotropic stress. This map was subsequently discretized into three categories (tensile regions, corresponding to averaged isotropic stresses larger than 100 Pa, compressive regions, corresponding to averaged isotropic stresses smaller than $-100$ Pa, and mixed regions, corresponding to intermediate values) and compared with the generated cell cycle S-G2-M duration map.

#### Plotting, statistics and reproducibility

All plotted heatmaps were smoothed using linear interpolation. All plots and graphs display the median of the corresponding distribution, unless stated otherwise. Similarly, all error bars represent 95% confidence intervals, unless otherwise specified. Statistical tests were conducted using MATLAB's Statistics and Machine Learning Toolbox, and $p$ values were reported up to four decimal places to take into account the significance. Probability Density Function (PDF) statistical distributions of any parameters were plotted using MATLAB's Statistics and Machine Learning Toolbox, using the function `histogram2` for 2D distribution, and `histcounts` for 1D distribution, with the normalization parameter set on pdf. All selected images correspond to quantified parameters and depict a representative state of the independent experiments reported. Unless stated otherwise, all images are representative of at least

$N = 2$ independent experiments. Sketches of proposed mechanisms and illustrations were created by hands as digital drawings using Clip Studio Paint and Inkscape.

#### Lyotropic active nematic model with tunable cell concentration

We developed a minimal continuum model that describes epithelia as a lyotropic active gel with tunable cell concentration ($\phi$) [22, 23]. The system is characterized by three key fields: the cell concentration $\phi$, the velocity field $\mathbf{u}$, and the nematic order parameter $\mathbf{Q} = 2q(\mathbf{nn} - \mathbf{I}/2)$, where $\mathbf{n}$ is the nematic director and $q$ is the magnitude of nematic order. The concentration $\phi$ varies from 0 (low density) to 1 (high density), representing local cell density.

The evolution of the nematic tensor is governed by

$$(\partial_t + \mathbf{u} \cdot \nabla)\mathbf{Q} - \mathbf{S} = \Gamma_Q \mathbf{H}, \tag{1}$$

where $\mathbf{S} = \lambda \mathbf{E} - (\boldsymbol{\omega} \cdot \mathbf{Q} - \mathbf{Q} \cdot \boldsymbol{\omega})$ is a generalized advection term, with $\mathbf{E} = (\nabla \mathbf{u} + \nabla \mathbf{u}^T)/2$ the strain rate tensor and $\boldsymbol{\omega} = (\nabla \mathbf{u}^T - \nabla \mathbf{u})/2$ the vorticity tensor. The alignment parameter $\lambda$ depends on particle shape: $\lambda > 0$ for rod-like particles, $\lambda < 0$ for disk-like particles, and $\lambda = 0$ for spherical particles. The molecular field $\mathbf{H}$ is defined as $\mathbf{H} = -\frac{\partial \mathcal{F}_{\mathrm{LC}}}{\partial \mathbf{Q}} + \nabla \cdot \left( \frac{\partial \mathcal{F}_{\mathrm{LC}}}{\partial (\nabla \mathbf{Q})} \right)$, where $\mathcal{F}_{\mathrm{LC}}$ represents the nematic elastic free energy based on the Oseen–Frank form under the single elastic constant approximation [24]: $\mathcal{F}_{\mathrm{LC}} = \frac{1}{2} K (\nabla \mathbf{Q})^2$.

Cell division is incorporated as a local source term in the equation for $\phi$. This formulation has been widely used to describe collective cell dynamics and reproduce experimentally observed proliferation and stress patterns [25, 26]. The dynamics of $\phi$ obey:

$$\partial_t \phi + \nabla \cdot (\mathbf{u}\phi) = \Gamma_\phi \nabla^2 \mu + \alpha \phi, \tag{2}$$

where $\Gamma_\phi$ is the mobility, $\alpha$ is the division rate, and the chemical potential is given by

$$\mu = \frac{\partial \mathcal{F}}{\partial \phi} - \nabla \cdot \left( \frac{\partial \mathcal{F}}{\partial \nabla \phi} \right). \tag{3}$$

The total free energy density $\mathcal{F}$ consists of both nematic and Ginzburg–Landau contributions:

$$\mathcal{F} = \mathcal{F}_{\mathrm{LC}} + \mathcal{F}_{\mathrm{GL}}, \tag{4}$$

where

$$\mathcal{F}_{\mathrm{GL}} = \frac{A_\phi}{2}\phi^2(1-\phi)^2 + \frac{K_\phi}{2}(\nabla\phi)^2, \tag{5}$$

describes phase separation and interfacial tension between isotropic and nematic regions [27].

The velocity field $\mathbf{u}$ evolves according to the momentum conservation equation:

$$\rho(\partial_t + \mathbf{u}\cdot\nabla)\mathbf{u} = \nabla \cdot \mathbf{\Pi}, \tag{6}$$

where $\rho$ is the effective mass density and $\mathbf{\Pi}$ is the total stress tensor, composed of active, viscous, elastic, and capillary contributions. The active stress, capturing contractility-driven nematic interactions, is modeled as:

$$\mathbf{\Pi}^{\mathrm{active}} = \zeta(1-\phi)\mathbf{Q}, \tag{7}$$

where $\zeta$ is the activity coefficient. This form reflects the experimentally observed suppression of activity with increasing cell density (Fig. S3C,D). The viscous stress is given by:

$$\mathbf{\Pi}^{\mathrm{viscous}} = 2\eta\mathbf{E}, \tag{8}$$

with $\eta$ the shear viscosity. The elastic stress is expressed as:

$$\mathbf{\Pi}^{\text{elastic}} = -P\mathbf{I} - \lambda\mathbf{H} + \mathbf{Q}\cdot\mathbf{H} - \mathbf{H}\cdot\mathbf{Q} - \nabla\mathbf{Q} : \frac{\partial\mathcal{F}}{\partial\nabla\mathbf{Q}}, \tag{9}$$

where $P$ is the reference pressure, set to zero in this study. The capillary stress accounts for chemical potential gradients and interfacial effects:

$$\mathbf{\Pi}^{\text{cap}} = (\mathcal{F} - \mu\phi)\mathbf{I} - \nabla\phi\left(\frac{\partial\mathcal{F}}{\partial\nabla\phi}\right), \tag{10}$$

where the first term represents a bulk pressure arising from crowding-induced increases in intracellular osmotic pressure, as reported in [28].

We use a hybrid Lattice Boltzmann algorithm to solve the coupled equations of motion. Full details of the numerical implementation are available in previous studies [29, 30]. Unless otherwise specified, simulations are performed using the following parameter values (in lattice units): $\Gamma_Q = 0.3$, $\lambda = 0.3$, $K = 0.002$, $\Gamma_\phi = 0.05$, $\zeta = 0.02$, $A_\phi = 0.03$, $K_\phi = 0.06$, and $\eta = 2/3$. For simulations with uniform cell concentration, we set $\alpha = 0$ and gradually increased $\phi$ from 0.3 to 0.7. For simulations with localized division, the initial cell concentration was set to $\phi = 0.5$, and division events were triggered at sites of high anisotropic stress, defined within a radius of 3 lattice units around the local maximum of anisotropic stress. Division events were introduced every $t_d = 150$ time steps and persisted for $\Delta t = 100$ time steps. The simulation domain was set to $200 \times 200$ lattice units and run for a total of 20,000 time steps. All results were computed from the final 3,000 steps to ensure statistical steady-state.

### Supporting Information

#### Invariants of the 2D stress tensor

The stress tensor $\boldsymbol{\sigma} = \begin{bmatrix} \sigma_{xx} & \sigma_{xy} \\ \sigma_{yx} & \sigma_{yy} \end{bmatrix}$ considered in this paper is treated as the 2D Cauchy stress tensor of continuum mechanics. The Cauchy stress tensor is symmetrical ($\sigma_{xy} = \sigma_{yx}$), and can be expressed in any system of coordinates using basis orientation transformation : $\boldsymbol{\sigma'} = \mathrm{D} \cdot \boldsymbol{\sigma} \cdot \mathrm{D}^T$ with $\mathrm{D} = \begin{bmatrix} \cos\theta & \sin\theta \\ -\sin\theta & \cos\theta \end{bmatrix}$, $\theta$ being the angle between the two systems of coordinates (x, y) and (x', y'). There are invariants associated with the stress tensor whose values do not depend on the coordinate system chosen. For the 2D stress tensor and using the Cayley–Hamilton theorem of linear algebra, first and second invariants can be defined as:

$$\mathrm{I}_1 = \mathrm{Tr}(\boldsymbol{\sigma}) = \sigma_{xx} + \sigma_{yy} \tag{11}$$

$$\mathrm{I}_2 = \det(\sigma) = \sigma_{xx}\sigma_{yy} - \sigma_{xy}^2 \tag{12}$$

In this paper, we consider the first invariant of the stress tensor $\mathrm{I}_1$. $\mathrm{I}_1$ quantifies the effect of mean normal stress at the point of space where the tensor is measured, therefore we define the isotropic stress as $\sigma_{iso} = \frac{\mathrm{I}_1}{2} = \frac{\sigma_{xx}+\sigma_{yy}}{2}$. $\sigma_{iso} > 0$ represents a case of local tension and $\sigma_{iso} < 0$ represents a case of local compression. Following the framework of continuum mechanics, $\boldsymbol{\sigma}$ can also be formalized as the sum of two sub-tensors:

$$\boldsymbol{\sigma} = \sigma_{iso}\mathbf{Id}_2 + \boldsymbol{\sigma_s}$$

$\boldsymbol{\sigma_s}$ is called the deviatoric stress tensor, and accounts for the non-isotropic part of the stress tensor. For the 2D deviator stress tensor, since $\boldsymbol{\sigma_s} = \boldsymbol{\sigma} - \sigma_{iso}\mathbf{Id}_2 = \begin{bmatrix} \frac{1}{2}(\sigma_{xx} - \sigma_{yy}) & \sigma_{xy} \\ \sigma_{yx} & \frac{1}{2}(\sigma_{yy} - \sigma_{xx}) \end{bmatrix}$, the first and second invariants can be expressed as:

$$\mathrm{J}_1 = \mathrm{Tr}(\boldsymbol{\sigma_s}) = 0 \tag{13}$$

$$\mathrm{J}_2 = -\det(\sigma_s) = [(\frac{\sigma_{xx} - \sigma_{yy}}{2})^2 + \sigma_{xy}^2] \tag{14}$$

One might observe that we compute $J_2$ with a minus sign comparing to $I_2$ but because $\sigma_s$ is traceless, both $\det(\sigma_s)$ and $-\det(\sigma_s)$ are solution of the characteristic polynomial. As we want a solution with a physical meaning (in term of tensor norm), we choose the positive one. In this paper, we consider the second invariant of the deviatoric stress tensor : since $\sigma_s$ is in a state of pure shear, $\mathrm{J}_2$ quantifies the magnitude of the maximum shear applied. Because the dimension of $\mathrm{J}_2$ is one that is a stress to the power of two, it is often convenient to define a second quantity $\mu$ by setting $\mu = \sqrt{\mathrm{J}_2}$. $\mu$ is sometimes referred to as maximum shear stress [31] and as deviatoric stress [32]: in this paper, we decide to call it anisotropic stress $\sigma_{aniso}$, following the convention of separating $\boldsymbol{\sigma}$ into two sub-components, one isotropic and one anisotropic. Even if $\sigma_{iso}$ and $\sigma_{aniso}$ are invariants, their expressions is still depending on $\sigma_{xx}$, $\sigma_{xy}$ and $\sigma_{yy}$: in this paper, we choose to do our analysis in the coordinate system where $\sigma_{xy} = 0$. This can be simply achieved by diagonalizing $\boldsymbol{\sigma}$: we note $\boldsymbol{\sigma_d}$ the stress tensor equivalent to $\boldsymbol{\sigma}$ set in the basis where the shear stress is null. $\boldsymbol{\sigma_d}$ verifies:

$$\boldsymbol{\sigma_d} = \begin{bmatrix} \sigma_M & 0 \\ 0 & \sigma_m \end{bmatrix} \tag{15}$$

$\sigma_M$ and $\sigma_m$ are the two eigenvalues of $\boldsymbol{\sigma}$: they define two perpendicular orientations in which the shear stress is maximum and minimum. $\sigma_M$ is called the maximum principle stress and $\sigma_m$ is called the minimum principal stress. By construction, $\sigma_M > \sigma_m$ so the direction of application of $\sigma_M$ is always the direction of the maximum shear stress (Fig. S3A). In this paper, what we call the principal stress direction is the orientation of application of $\sigma_M$. Using the maximum and minimum principal stress, $\sigma_{iso}$ and $\sigma_{aniso}$ become:

$$\sigma_{iso} = \frac{\sigma_M + \sigma_m}{2} \tag{16}$$

$$\sigma_{aniso} = \frac{\sigma_M - \sigma_m}{2} \tag{17}$$

#### Stress topological defects

Topological defects are a universal class of spatial patterns found in multiple fields of physics [18, 33, 34]. Simply put, they can be defined as regions where a given parameter field collapses locally and can no longer be defined : this constitutes a topological singularity, or a topological defect. In this work, the existence of topological stress defects arises from the definition of the principal stress direction. In 2D, the relationship between $\boldsymbol{\sigma_d}$ and $\boldsymbol{\sigma}$ involves reorienting $\boldsymbol{\sigma}$ in a basis with orientation $\theta_p$, where $\theta_p$ is the principal stress direction. We have: $\boldsymbol{\sigma_d} = \mathrm{D_p} \cdot \boldsymbol{\sigma} \cdot \mathrm{D_p}^T$ with $\mathrm{D_p} = \begin{bmatrix} \cos\theta_p & \sin\theta_p \\ -\sin\theta_p & \cos\theta_p \end{bmatrix}$ Projecting the equation, we have the system:

$$\begin{cases} \sigma_M = \sigma_{xx}\cos^2\theta_p + \sigma_{yy}\sin^2\theta_p + 2\sigma_{xy}\sin\theta_p\cos\theta_p \\ \sigma_m = \sigma_{xx}\sin^2\theta_p + \sigma_{yy}\cos^2\theta_p - 2\sigma_{xy}\sin\theta_p\cos\theta_p \\ 0 = (\sigma_{yy} - \sigma_{xx})\sin\theta_p\cos\theta_p + \sigma_{xy}(\cos^2\theta_p - \sin^2\theta_p) \end{cases}$$

This system of equation is usefull to express $\sigma_M$, $\sigma_m$ and $\theta_p$ only using a function of $(\sigma_{xx}, \sigma_{yy}, \sigma_{xy})$. In particular, the expression concerning $\theta_p$ is :

$$\tan 2\theta_p = \frac{2\sigma_{xy}}{(\sigma_{xx} - \sigma_{yy})} \tag{18}$$

From here, one can see directly that this equation is not defined when $(\sigma_{xx} - \sigma_{yy)} \rightarrow 0$. By definition, any local region where this condition is approached constitutes a topological defect, as $\theta_p$ becomes undefined. Since this property holds in any basis, it is natural that topological defects theoretically emerge in regions where $\sigma_{\text{aniso}} \rightarrow 0$.

It is important to emphasize that in this paper, we do not define stress topological defects based on this analytical condition. Instead, we compute the topological charge using a winding number approach (see Methods). Through this method, we experimentally recover the natural mathematical property expressed in equation 18.

### Supplementary Table

**Table 1 A general mechanical principle for cell division across model system**. This table includes studies reporting cell division alignment with anisotropic stress-based cues, unless otherwise specified. When possible, we also report comparisons with cell-shape–based cues to ensure consistent evaluation across studies.

| Author | System | Stress | Measure of stress | Source of stress | Shape cue |
|---|---|---|---|---|---|
| Lintilhac et al. (1984) [35] | Nicotiana tabacum pith tissue [1] | compressive | ∅ | **Ext**, Uniaxial compression | NA |
| Minc et al. (2011) [36] | Sea urchin zygote | compressive | ∅ | **Ext**, anisotropic CS | yes |
| Minc et al. (2011) [36] | Sea urchin zygote | compressive[2] | ∅ | **Ext**, isotropic CS | × |
| Fink et al. (2011) [37] | HeLa/RPE1 single cells | tensile | Laser ablation | **Ext**, anisotropic BC | yes |
| Fink et al. (2011) [37] | HeLa/RPE1 single cells | tensile[3] | Laser ablation | **Ext**, isotropic BC | × |
| LeGoff et al. (2013) [38] | Drosophila wing disc | tensile | Laser ablation | Self-organized | yes |
| Campinho et al. (2013) [39] | Zebrafish epiboly | tensile | Laser ablation | Self-organized | yes |
| Wyatt et al. (2015) [40] | MDCK suspended monolayer | tensile | Laser ablation | **Ext**, Uniaxial stretching | yes |
| Wyatt et al. (2015) [40] | MDCK suspended monolayer [*subset*] | tensile [4] | Laser ablation | **Ext**, Uniaxial stretching | yes |
| Bosveld et al. (2016) [41] | Drosophila pupal notum | mixed [5] | Laser ablation | Self-organized | yes |
| Louveaux et al. (2016) [42] | Arabidopsis thaliana shoot apex [1] | tensile | Laser ablation | Self-organized | NA |
| Hart et al. (2017) [43] | MDCK monolayer | tensile | ∅ | **Ext**, Uniaxial stretching | × |
| Finegan et al. (2018) [44] | Drosophila follicular epithelium | tensile | Laser ablation | Self-organized | × |
| Scarpa et al. (2018) [45] | Drosophila embryo, cell cycle 9 to 11 | tensile | Laser ablation | Self-organized | × |
| Nestor-Bergmann et al. (2019) [46] | Xenopus tissue embryo | tensile | ∅ | **Ext**, Uniaxial stretching | yes |
| Nestor-Bergmann et al. (2019) [46] | Xenopus tissue embryo | mixed [6] | Vertex-based model | **Ext**, Uniaxial stretching | yes |
| Lisica et al. (2022) [47] | MDCK suspended monolayer | tensile | ∅ | **Ext**, Uniaxial stretching | yes |
| Lisica et al. (2022) [47] | MDCK suspended monolayer | compressive [7] | ∅ | **Ext**, Uniaxial compression | × |
| Blanchard et al. (2024) [48] | Drosophila embryo, cell cycle 14 | compressive | Laser ablation | Self-organized | × [8] |
| Middelkoop et al. (2024) [49] | C. elegans + mouse zygote | compressive | ∅ | **Ext**, CS | yes |

Footnotes appear on the following page.

∅ : not reported or not assessed in the original study. When the mechanical state was not assessed in the original study, we inferred it qualitatively based on the source of stress.

**Ext** : the internal stress organization is imposed by external constraints.

NA : Not applicable.

BC : Boundary conditions enforcing stress organization because of micropatterning.

CS : Compressive shell (or for Minc et al. micro-fabricated chambers).

[1]These studies were performed in plant epidermis rather than animal epithelia, but we considered them relevant because both systems share important features, including their organization as densely packed cellular sheets, and the existence of tissue-scale mechanical coupling. We chose to include them in this table because they report qualitatively similar couplings between anisotropic stress and division orientation, suggesting that the relationship identified here may extend beyond the specific epithelial contexts considered in this study.

[2]Uniaxial (z-axis) compressive confinement with isotropic in-plane geometry. The authors report cell division randomly oriented within the xy-plane but restricted to the plane perpendicular to compression.

[3]Isotropic tension generated by an isotropic pattern. Cells exhibit random in-plane division orientation under these conditions. Interestingly, external stretching of the pattern restores alignment of division orientation.

[4]In the subset (5%) of cells not aligning their division axis with the stretch direction, the authors show that cell long axis remains predictive of division orientation.

[5]The authors performed large-scale laser ablation and observed anisotropic recoil of the surrounding tissue.

[6]The authors report that division orientation is not determined by the global stretch orientation but correlates with local stress anisotropy and cell shape anisotropy.

[7]The authors report that under compressive loading, epithelial monolayers exhibit an increased fraction of out-of-plane divisions. This orientation is consistent with the axis of lowest compressive stress in a suspended monolayer and should correspond to maximal stress anisotropy direction.

[8]The authors report that high stress magnitude can override initial cell shape anisotropy.

### References


[1] Balasubramaniam, L., Jain, S., Dang, T., Lagoutte, E., Marc Mège, R., Chavrier, P., Ladoux, B., Rossé, C.: Different biomechanical cell behaviors in an epithelium drive collective epithelial cell extrusion. Advanced Science, 2401573 (2024)

[2] Streichan, S.J., Hoerner, C.R., Schneidt, T., Holzer, D., Hufnagel, L.: Spatial constraints control cell proliferation in tissues. Proceedings of the National Academy of Sciences **111**(15), 5586–5591 (2014)

[3] Dickson, M.A., Hahn, W.C., Ino, Y., Ronfard, V., Wu, J.Y., Weinberg, R.A., Louis, D.N., Li, F.P., Rheinwald, J.G.: Human keratinocytes that express htert and also bypass a p16ink4a-enforced mechanism that limits life span become immortal yet retain normal growth and differentiation characteristics. Molecular and cellular biology **20**(4), 1436–1447 (2000)

[4] Xi, W., Saleh, J., Yamada, A., Tomba, C., Mercier, B., Janel, S., Dang, T., Soleilhac, M., Djemat, A., Wu, H., *et al.*: Modulation of designer biomimetic matrices for optimized differentiated intestinal epithelial cultures. Biomaterials **282**, 121380 (2022)

[5] Barai, A., Soleilhac, M., Xi, W., Lin, S.-Z., Karnat, M., Bazellières, E., Richelme, S., Lecouffe, B., Chardès, C., Berrebi, D., *et al.*: A multicellular star-shaped actin network underpins epithelial organization and connectivity. Nature Communications **16**(1), 6201 (2025)

[6] Abe, T., Kiyonari, H., Shioi, G., Inoue, K.-I., Nakao, K., Aizawa, S., Fujimori, T.: Establishment of conditional reporter mouse lines at rosa26 locus for live cell imaging. genesis **49**(7), 579–590 (2011)

[7] Krndija, D., El Marjou, F., Guirao, B., Richon, S., Leroy, O., Bellaiche, Y., Hannezo, E., Matic Vignjevic, D.: Active cell migration is critical for steady-state epithelial turnover in the gut. Science **365**(6454), 705–710 (2019)

[8] Pérez-González, C., Ceada, G., Greco, F., Matejčić, M., Gómez-González, M., Castro, N., Menendez, A., Kale, S., Krndija, D., Clark, A.G., *et al.*: Mechanical compartmentalization of the intestinal organoid enables crypt folding and collective cell migration. Nature cell biology **23**(7), 745–757 (2021)

[9] Schoenit, A., Monfared, S., Anger, L., Rosse, C., Venkatesh, V., Balasubramaniam, L., Marangoni, E., Chavrier, P., Mège, R.-M., Doostmohammadi, A., et al.: Force transmission is a master regulator of mechanical cell competition. Nature Materials, 1–11 (2025)

[10] Schindelin, J., Arganda-Carreras, I., Frise, E., Kaynig, V., Longair, M., Pietzsch, T., Preibisch, S., Rueden, C., Saalfeld, S., Schmid, B., *et al.*: Fiji: an open-source platform for biological-image analysis. Nature methods **9**(7), 676–682 (2012)

[11] Stringer, C., Pachitariu, M.: Cellpose3: one-click image restoration for improved cellular segmentation. Nature Methods, 1–8 (2025)

[12] Tinevez, J.-Y., Perry, N., Schindelin, J., Hoopes, G.M., Reynolds, G.D., Laplantine, E., Bednarek, S.Y., Shorte, S.L., Eliceiri, K.W.: Trackmate: An open and extensible platform for single-particle tracking. Methods **115**, 80–90 (2017)

[13] Thielicke, W., Sonntag, R.: Particle image velocimetry for MATLAB: Accuracy and enhanced algorithms in PIVlab. Journal of Open Research Software **9**(1), 12 (2021) https://doi.org/10.5334/jors.334

[14] Püspöki, Z., Storath, M., Sage, D., Unser, M.: Transforms and operators for directional bioimage analysis: a survey. Focus on bio-image informatics, 69–93

(2016)

[15] Monfared, S., Ravichandran, G., Andrade, J., Doostmohammadi, A.: Mechanical basis and topological routes to cell elimination. Elife **12**, 82435 (2023)

[16] Saw, T.B., Doostmohammadi, A., Nier, V., Kocgozlu, L., Thampi, S., Toyama, Y., Marcq, P., Lim, C.T., Yeomans, J.M., Ladoux, B.: Topological defects in epithelia govern cell death and extrusion. Nature **544**(7649), 212–216 (2017)

[17] Balasubramaniam, L., Doostmohammadi, A., Saw, T.B., Narayana, G.H.N.S., Mueller, R., Dang, T., Thomas, M., Gupta, S., Sonam, S., Yap, A.S., *et al.*: Investigating the nature of active forces in tissues reveals how contractile cells can form extensile monolayers. Nature materials **20**(8), 1156–1166 (2021)

[18] Vromans, A.J., Giomi, L.: Orientational properties of nematic disclinations. Soft matter **12**(30), 6490–6495 (2016)

[19] Li, K.: The image stabilizer plugin for ImageJ. http://www.cs.cmu.edu/~kangli/code/Image_Stabilizer.html (2008)

[20] Nier, V., Jain, S., Lim, C.T., Ishihara, S., Ladoux, B., Marcq, P.: Inference of internal stress in a cell monolayer. Biophysical journal **110**(7), 1625–1635 (2016)

[21] Peyret, G., Mueller, R., d'Alessandro, J., Begnaud, S., Marcq, P., Mège, R.-M., Yeomans, J.M., Doostmohammadi, A., Ladoux, B.: Sustained oscillations of epithelial cell sheets. Biophysical journal **117**(3), 464–478 (2019)

[22] Thampi, S.P., Doostmohammadi, A., Golestanian, R., Yeomans, J.M.: Intrinsic free energy in active nematics. Europhysics Letters **112**(2), 28004 (2015)

[23] Doostmohammadi, A., Thampi, S.P., Yeomans, J.M.: Defect-mediated morphologies in growing cell colonies. Physical review letters **117**(4), 048102 (2016)

[24] De Gennes, P.-G., Prost, J.: The Physics of Liquid Crystals vol. 83. Oxford university press, ??? (1993)

[25] Rossen, N.S., Tarp, J.M., Mathiesen, J., Jensen, M.H., Oddershede, L.B.: Long-range ordered vorticity patterns in living tissue induced by cell division. Nature communications **5**(1), 5720 (2014)

[26] Doostmohammadi, A., Thampi, S.P., Saw, T.B., Lim, C.T., Ladoux, B., Yeomans, J.M.: Celebrating soft matter's 10th anniversary: Cell division: a source of active stress in cellular monolayers. Soft Matter **11**(37), 7328–7336 (2015)

[27] Cahn, J.W., Hilliard, J.E.: Free energy of a nonuniform system. i. interfacial free energy. The Journal of chemical physics **28**(2), 258–267 (1958)

[28] Vian, A., Pochitaloff, M., Yen, S.-T., Kim, S., Pollock, J., Liu, Y., Sletten, E.M., Campàs, O.: In situ quantification of osmotic pressure within living embryonic tissues. Nature Communications **14**(1), 7023 (2023)

[29] Marenduzzo, D., Orlandini, E., Cates, M., Yeomans, J.: Steady-state hydrodynamic instabilities of active liquid crystals: Hybrid lattice boltzmann simulations. Physical Review E—Statistical, Nonlinear, and Soft Matter Physics **76**(3), 031921 (2007)

[30] Denniston, C., Marenduzzo, D., Orlandini, E., Yeomans, J.: Lattice boltzmann algorithm for three–dimensional liquid–crystal hydrodynamics. Philosophical Transactions of the Royal Society of London. Series A: Mathematical, Physical and Engineering Sciences **362**(1821), 1745–1754 (2004)

[31] Tambe, D.T., Corey Hardin, C., Angelini, T.E., Rajendran, K., Park, C.Y., Serra-Picamal, X., Zhou, E.H., Zaman, M.H., Butler, J.P., Weitz, D.A., *et al.*: Collective cell guidance by cooperative intercellular forces. Nature materials **10**(6), 469–475

(2011)

[32] Gauquelin, E., Kuromiya, K., Namba, T., Ikawa, K., Fujita, Y., Ishihara, S., Sugimura, K.: Mechanical convergence in mixed populations of mammalian epithelial cells. The European Physical Journal E **47**(3), 21 (2024)

[33] Fumeron, S., Berche, B.: Introduction to topological defects: from liquid crystals to particle physics. The European Physical Journal Special Topics **232**(11), 1813–1833 (2023)

[34] Fardin, M.-A., Ladoux, B.: Living proof of effective defects. Nature Physics **17**(2), 172–173 (2021)

[35] Lintilhac, P.M., Vesecky, T.B.: Stress-induced alignment of division plane in plant tissues grown in vitro. Nature **307**(5949), 363–364 (1984)

[36] Minc, N., Burgess, D., Chang, F.: Influence of cell geometry on division-plane positioning. Cell **144**(3), 414–426 (2011)

[37] Fink, J., Carpi, N., Betz, T., Bétard, A., Chebah, M., Azioune, A., Bornens, M., Sykes, C., Fetler, L., Cuvelier, D., *et al.*: External forces control mitotic spindle positioning. Nature cell biology **13**(7), 771–778 (2011)

[38] LeGoff, L., Rouault, H., Lecuit, T.: A global pattern of mechanical stress polarizes cell divisions and cell shape in the growing drosophila wing disc. Development **140**(19), 4051–4059 (2013)

[39] Campinho, P., Behrndt, M., Ranft, J., Risler, T., Minc, N., Heisenberg, C.-P.: Tension-oriented cell divisions limit anisotropic tissue tension in epithelial spreading during zebrafish epiboly. Nature cell biology **15**(12), 1405–1414 (2013)

[40] Wyatt, T.P., Harris, A.R., Lam, M., Cheng, Q., Bellis, J., Dimitracopoulos, A.,

Kabla, A.J., Charras, G.T., Baum, B.: Emergence of homeostatic epithelial packing and stress dissipation through divisions oriented along the long cell axis. Proceedings of the National Academy of Sciences **112**(18), 5726–5731 (2015)

[41] Bosveld, F., Markova, O., Guirao, B., Martin, C., Wang, Z., Pierre, A., Balakireva, M., Gaugue, I., Ainslie, A., Christophorou, N., *et al.*: Epithelial tricellular junctions act as interphase cell shape sensors to orient mitosis. Nature **530**(7591), 495–498 (2016)

[42] Louveaux, M., Julien, J.-D., Mirabet, V., Boudaoud, A., Hamant, O.: Cell division plane orientation based on tensile stress in arabidopsis thaliana. Proceedings of the National Academy of Sciences **113**(30), 4294–4303 (2016)

[43] Hart, K.C., Tan, J., Siemers, K.A., Sim, J.Y., Pruitt, B.L., Nelson, W.J., Gloerich, M.: E-cadherin and lgn align epithelial cell divisions with tissue tension independently of cell shape. Proceedings of the National Academy of Sciences **114**(29), 5845–5853 (2017)

[44] Finegan, T.M., Na, D., Cammarota, C., Skeeters, A.V., Nádasi, T.J., Dawney, N.S., Fletcher, A.G., Oakes, P.W., Bergstralh, D.T.: Tissue tension and not interphase cell shape determines cell division orientation in the drosophila follicular epithelium. The EMBO journal **38**(3), 100072 (2019)

[45] Scarpa, E., Finet, C., Blanchard, G.B., Sanson, B.: Actomyosin-driven tension at compartmental boundaries orients cell division independently of cell geometry in vivo. Developmental cell **47**(6), 727–740 (2018)

[46] Nestor-Bergmann, A., Stooke-Vaughan, G.A., Goddard, G.K., Starborg, T., Jensen, O.E., Woolner, S.: Decoupling the Roles of Cell Shape and Mechanical Stress in Orienting and Cueing Epithelial Mitosis. Cell Reports **26**(8), 2088–21004

(2019)

[47] Lisica, A., Fouchard, J., Kelkar, M., Wyatt, T.P., Duque, J., Ndiaye, A.-B., Bonfanti, A., Baum, B., Kabla, A.J., Charras, G.T.: Tension at intercellular junctions is necessary for accurate orientation of cell division in the epithelium plane. Proceedings of the National Academy of Sciences **119**(49), 2201600119 (2022)

[48] Blanchard, G.B., Scarpa, E., Muresan, L., Sanson, B.: Mechanical stress combines with planar polarised patterning during metaphase to orient embryonic epithelial cell divisions. Development **151**(10), 202862 (2024)

[49] Middelkoop, T.C., Neipel, J., Cornell, C.E., Naumann, R., Pimpale, L.G., Jülicher, F., Grill, S.W.: A cytokinetic ring-driven cell rotation achieves hertwig's rule in early development. Proceedings of the National Academy of Sciences **121**(25), 2318838121 (2024)

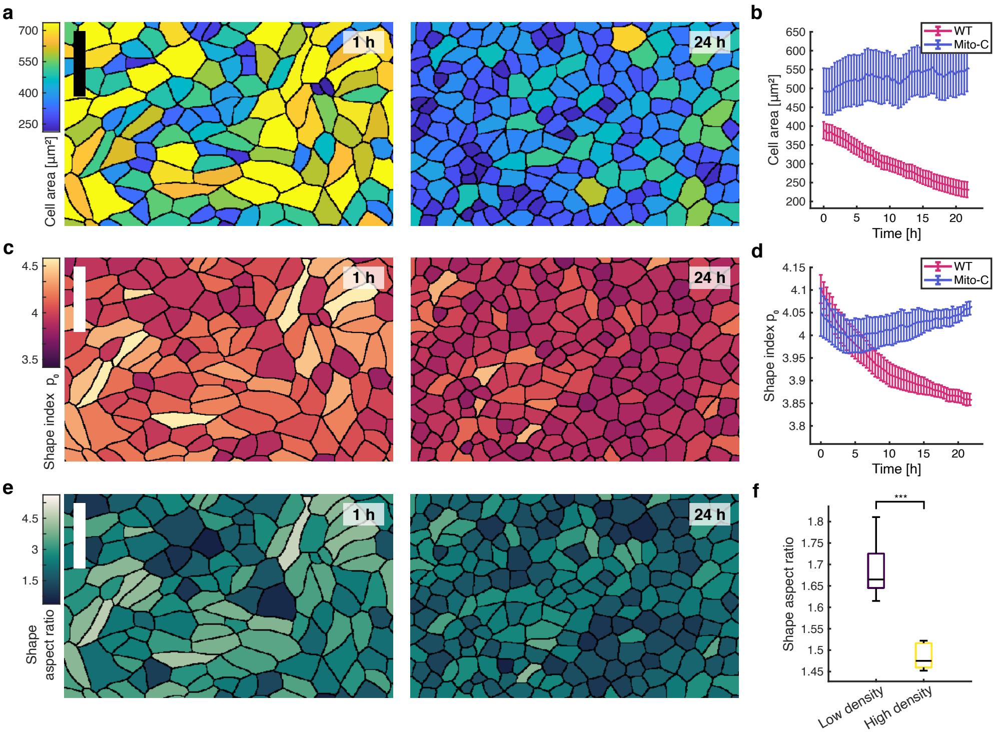


**Fig. S1 MDCK monolayer mechanical state evolves with tissue densification**. **a**) Left, segmented image of sub-confluent MDCK monolayer (t = 1h) at low density, right, same monolayer in high density (t = 24h). Color code is the cell area. **b**) Average (mean) evolution in time of the cell area within the whole tissue, for MDCK WT and for mitomycin-C treated MDCK cells. n = 15 movies from N = 3 independent experiments (WT) ; n = 15 movies from N = 3 independent experiments (Mito-C). **c**) Left, segmented image of sub-confluent MDCK monolayer (t = 1h) at low density, right, same monolayer in high density (t = 24h). Color code is the cell shape index. **d**) Average (mean) evolution in time of the cell shape index within the whole tissue, for MDCK WT and for mitomycin-C treated MDCK cells. n = 15 movies from N = 3 independent experiments (WT) ; n = 15 movies from N = 3 independent experiments (Mito-C). **e**) Left, segmented image of sub-confluent MDCK monolayer (t = 1h) at low density, right, same monolayer in high density (t = 24h). Color code is the cell aspect ratio. **f**) Average shape aspect ratio for low density (t = 1h) and high density (t = 24h). 45 points, n = 15 movies from N = 3 independent experiments. $p \leq 0.001$. [Scale bar 50 μm (a, c, e). P-value from Mann-Whitney U-test (f).]

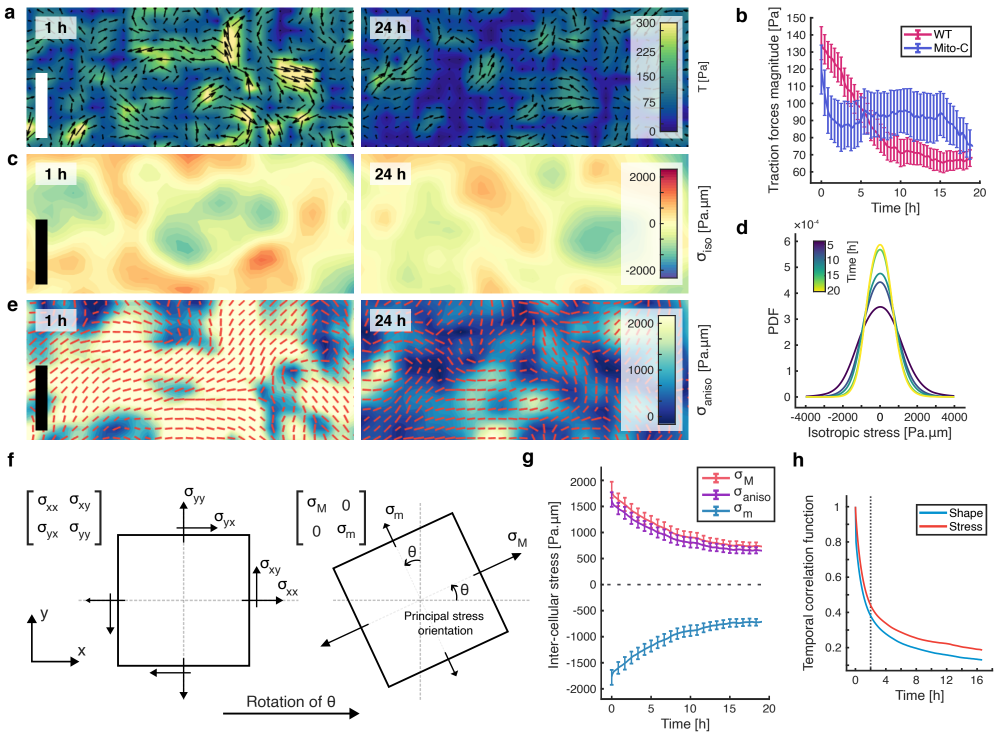


**Fig. S2** **MDCK monolayer mechanical state is anisotropic in nature**. **a**) Left, traction force field of sub-confluent MDCK monolayer (t = 1h) at low density, right, same field in high density (t = 24h). Color code is the traction force magnitude. **b**) Average evolution in time of the traction force magnitude within the whole tissue, for MDCK WT and for mitomycin-C treated MDCK cells. n = 24 movies from N = 5 independent experiments (WT) ; n = 15 movies from N = 3 independent experiments (Mito-C). **c**) Left, isotropic stress field of MDCK monolayer in a low density state (t = 1h), right, same field in high density state (t = 24h). Color code is the isotropic stress value. **d**) Probability density function (PDF) evolution of the isotropic stress distribution in MDCK monolayer during monolayer densification. Color code is the time. n = 24 movies from N = 5 independent experiments. **e**) Left, anisotropic stress field of MDCK monolayer in a low density state (t = 1h), right, same field in high density state. Color code is the anisotropic stress magnitude. Principal stress orientation director field in red. **f**) Simplified representation of the transformation allowing the computation of the maximum and minimum stress and defining the principal stress orientation. **g**) Evolution of average inter-cellular stress components in time within the wole tissue (maximum, minimum and anisotropic stress). n = 24 movies from N = 5 independent experiments. **h**) Decay of the temporal correlation function for the principal stress orientation and cell shape orientation. n = 13 positions from N = 3 independent experiments. [Scale bar 70 μm (a, c, e). All error bars show the confidence interval at 95 %.]

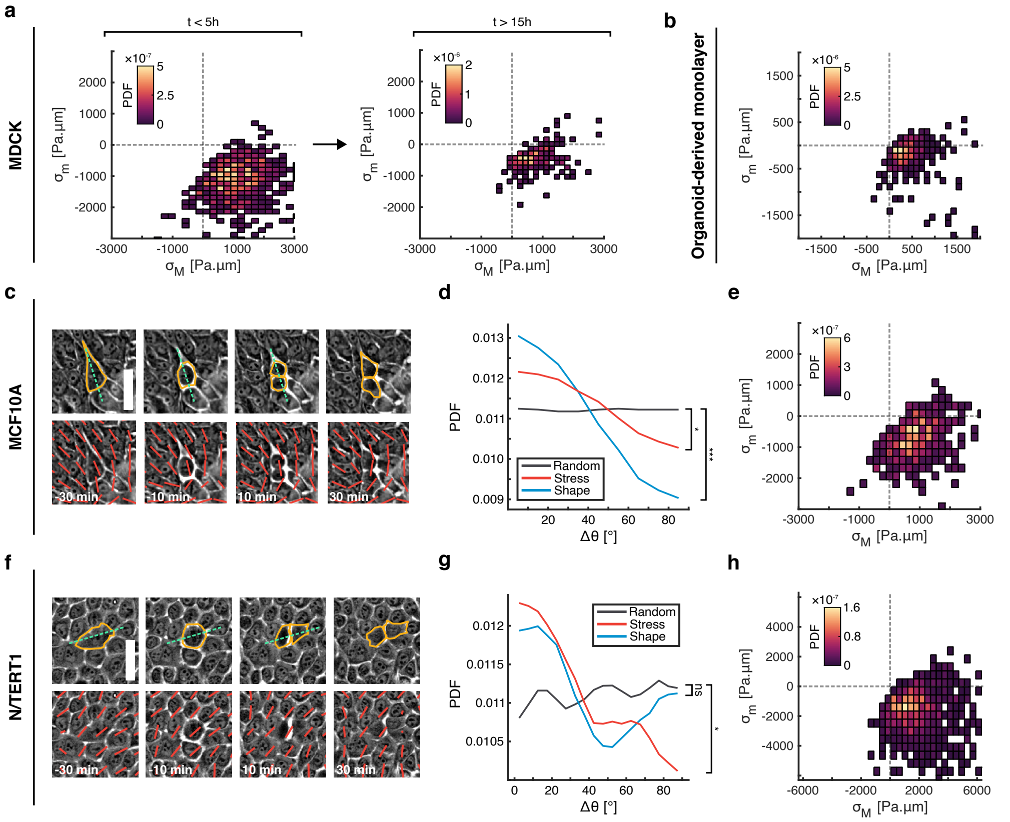


**Fig. S3 Principal stress orientation is a stable predictor of cell division orientation in epithelia**. **a**) 2D histogram of the maximum and minimum principal stress coupling 30 min before cytokinesis for MDCK monolayer, left, in low density state ($t < 5h$), and right, in high density state ($t > 15h$). Color code is the PDF. 767 cell divisions (($t < 5h$) and 150 cell divisions ($t > 15h$). $n = 13$ movies from $N = 2$ independent experiments. **b**) 2D histogram of the maximum and minimum principal stress coupling 30 min before cytokinesis for organoid-derived monolayer. Color code is the PDF. 526 divisions. $n = 13$ movies from $N = 3$ independent experiments. **c**) Phase contrast image of cell division process, showing in dotted white line cell division orientation, before and after cytokinesis for MCF10A monolayer. Principal stress orientation in red. **d**) Probability distribution function of the angle between cell division orientation and principal stress orientation, corresponding to (a). 354 cell divisions and 2348 random positions. $n = 7$ movies from $N = 2$ independent experiments. $p \leq 0.04$ (stress vs random), and $p \leq 0.001$ (shape vs random). **e**) 2D histogram of the maximum and minimum principal stress coupling 30 min before cytokinesis for MCF10A monolayer. Color code is the PDF. 354 divisions. $n = 7$ movies from $N = 2$ independent experiments. **f**) Phase contrast image of cell division process, showing in dotted white line cell division orientation, before and after cytokinesis for N/TERT1 monolayer. Principal stress orientation in red. **g**) Probability distribution function of the angle between cell division orientation and principal stress orientation, corresponding to (c). 850 cell divisions and 2100 random positions. $n = 7$ movies from $N = 3$ independent experiments. $p \leq 0.04$ (stress vs random), and $p = 0.4311$ (shape vs random). **h**) 2D histogram of the maximum and minimum principal stress coupling 30 min before cytokinesis for N/TERT1 monolayer. Color code is the PDF. 850 divisions. $n = 7$ movies from $N = 3$ independent experiments. [Scale bar 70 µm (c, f). P-value from Mann-Whitney U-test (d, g)]

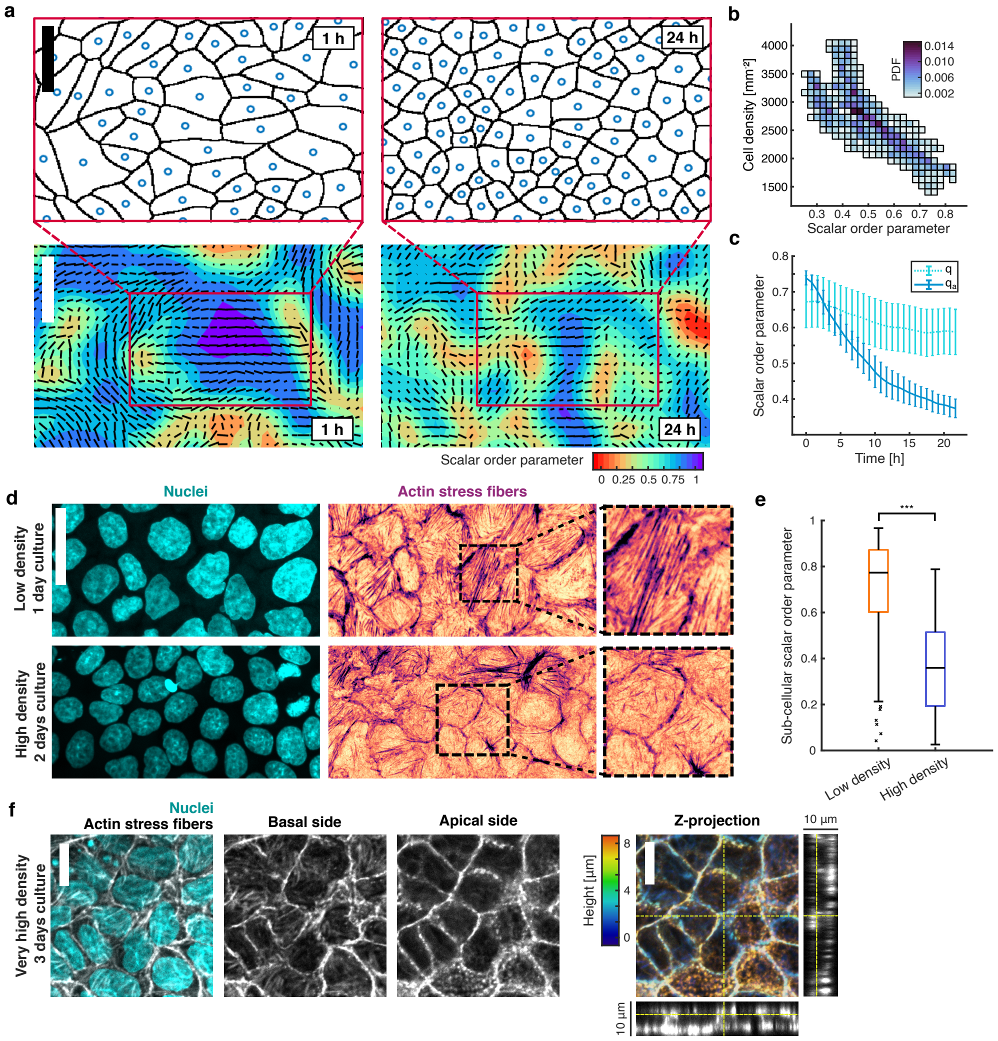


**Fig. S4 Tissue densification disrupts nematic order across multiple scales**. **a**) Top, left, segmented image of sub-confluent MDCK monolayer (t = 1h) at low density, right, same monolayer in high density (t = 24h). Blue dots represent cells center of mass. Bottom, left, scalar local order parameter for the nematic director of MDCK monolayer (t = 1h) at low density, right, same monolayer in high density (t = 24h). Color code is the local scalar order parameter. Nematic director field is in black. Magnitude of the elongation is weighed with the cell aspect ratio. The red box correspond to the segmented localization of the top panel. **b**) 2D histogram of the coupling between cell density and scalar order parameter across time. Color code is the probability density function (PDF). n = 15 movies from N = 3 independent experiments. **c**) Average scalar order parameter in time for the nematic director field, using direct ($q$) computation or normalized ($q_a$, shape index is used as weights) computation. Using $q$ is not enough to make the order collapse because cell orientations computed past t = 10h do not have a physical meaning anymore (see Materials and Methods). n = 15 movies from N = 3 independent experiments. **d**) Example images of the actin cytoskeleton basal organization (left) for low density monolayer and higher density monolayer state (top and bottom). Right is the corresponding nuclei. **e**) Average intra-cellular actin fiber scalar order parameter for low and high cell density. 75 cells (low), 87 cells (high), n = 14 images (low) and n = 13 images (high) from N = 3 independent experiments. **f**) Typical images of the actin cytoskeleton basal and apical organization at very high cell density, with the corresponding nuclei staining. Left is the corresponding z-projection, colored by the height of each slice. [Scale bar 75 µm (bottom a), 30 µm (top a, d), 10 µm (f). P-value from Mann-Whitney U-test (e).]

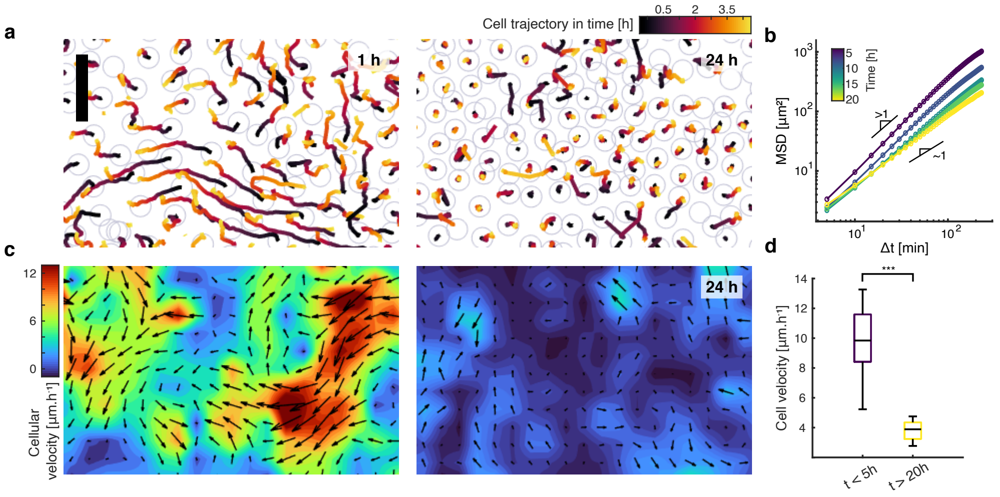


**Fig. S5 MDCK cells undergo jamming transition during tissue densification.** **a**) Cellular displacement trajectory during 4h for, left, a sub-confluent MDCK monolayer (t = 1h) in nematic state, and right, the same monolayer upon jamming transition (t = 24h). Color code is the time. **b**) Log-log mean square displacement (MSD) evolution during jamming transition. n = 15 movies from N = 3 independent experiments. Color code is the moment in time. **c**) Left, cellular velocity field of sub-confluent MDCK monolayer (t = 1h) in nematic state, right, same field upon jamming transition (t = 24h). Color code is the velocity magnitude. **d**) Average velocity magnitude during low and high cell density phase. 72 points, n = 24 movies from N = 5. $p \leq 0.001$. [Scale bar 40 µm (a, c). P-value from Mann-Whitney U-test (d).]

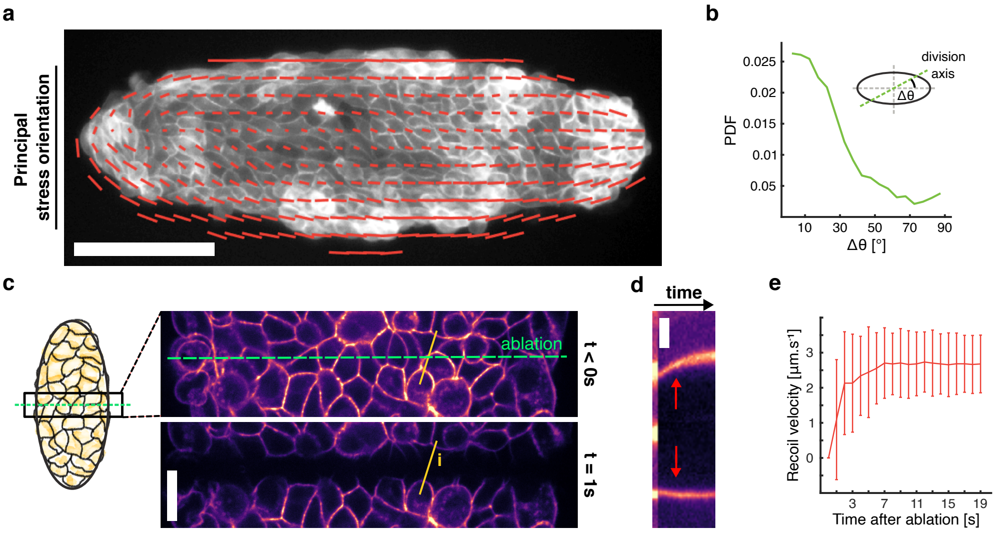


**Fig. S6 Patterning induced stress anisotropy controls cell division orientation**. **a**) CAAX GFP MDCK cells on ellipse shaped pattern. Principal stress orientation in red. **b**) Probability distribution function of the angle between cell division orientation and long axis of the ellipse. 228 cell divisions. n = 14 patterns from N = 2 independent experiments. **c**) CAAX GFP MDCK cells image focused on the center of the ellipse, right before and right after laser ablation across the small axis. **d**) Kymograph of cells temporal recoil after ablation, corresponding to the yellow line **i** in (c). **e**) Average (mean) evolution in time of the recoil velocity following laser ablation (happening at t = 0s). n = 14 patterns from N = 2 independent experiments. [Scale bar 125 µm (a), 15 µm (c), 2 µm (d).]

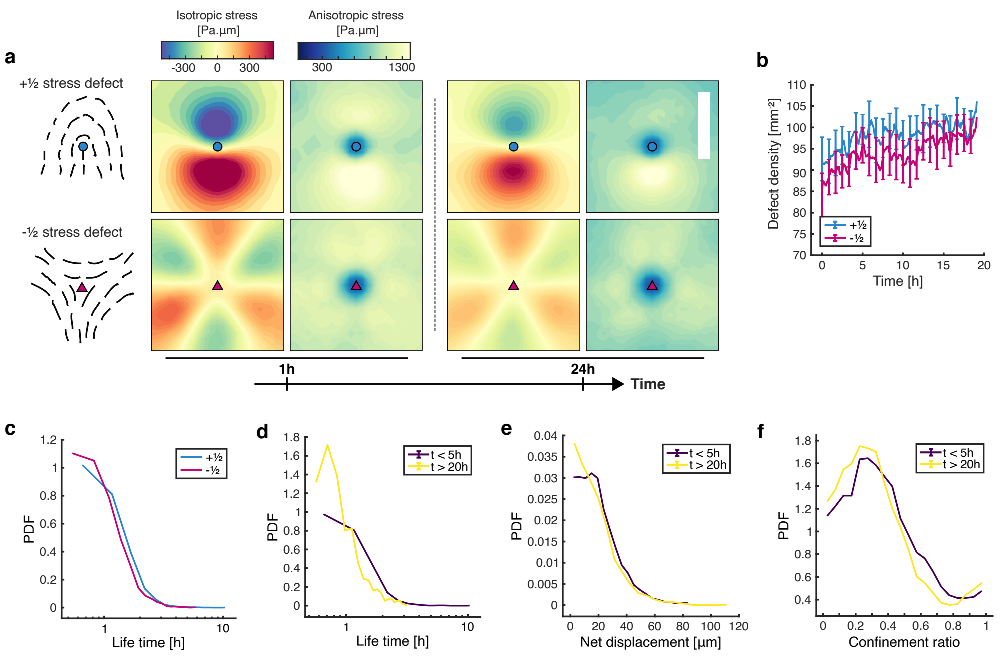


**Fig. S7 Topological stress defects behavior in space and time is stable during densification**. **a**) Isotropic (left) and anisotropic (right) stress field around stress topological defects oriented according to the schematic, during low and high density state. Top is $+\frac{1}{2}$ stress defects and bottom is $-\frac{1}{2}$ stress defects. Color code is the given stress component magnitude. 2441 $+\frac{1}{2}$ stress defects in low density state, 3152 $+\frac{1}{2}$ stress defects in high density state and 2346 $-\frac{1}{2}$ stress defects in low density state, 2894 $-\frac{1}{2}$ stress defects in high density state. n = 10 movies from N = 2 independent experiments. **b**) Average stress defects density evolution in time. n = 24 movies from N = 5 independent experiments. **c**) Probability distribution function (PDF) of one given stress defects life time in the tissue. n = 5 movies from N = 5 independent experiments. 2358 $+\frac{1}{2}$ stress defects and 1897 $-\frac{1}{2}$ stress defects. **d**) PDF of stress defects life time regardless of the charge for low and high density state. 839 stress defects tracked in the liquid phase and 644 stress defects tracked in the jammed phase. n = 5 movies from N = 5 independent experiments. **e**) PDF of stress defects net displacement regardless of the charge for low and high density state. 839 stress defects tracked in the liquid phase and 644 stress defects tracked in the jammed phase. n = 5 movies from N = 5 independent experiments. **f**) PDF of stress defects confinement ratio regardless of the charge for low and high density state. The confinement ratio ranges from 0 (confined or caged motion) to 1 (straight-line trajectory). 839 stress defects tracked in low density phase and 644 stress defects tracked in the high density phase. n = 5 movies from N = 5 independent experiments. [Scale bar 75 µm (a). All error bars show the confidence interval at 95 %.]

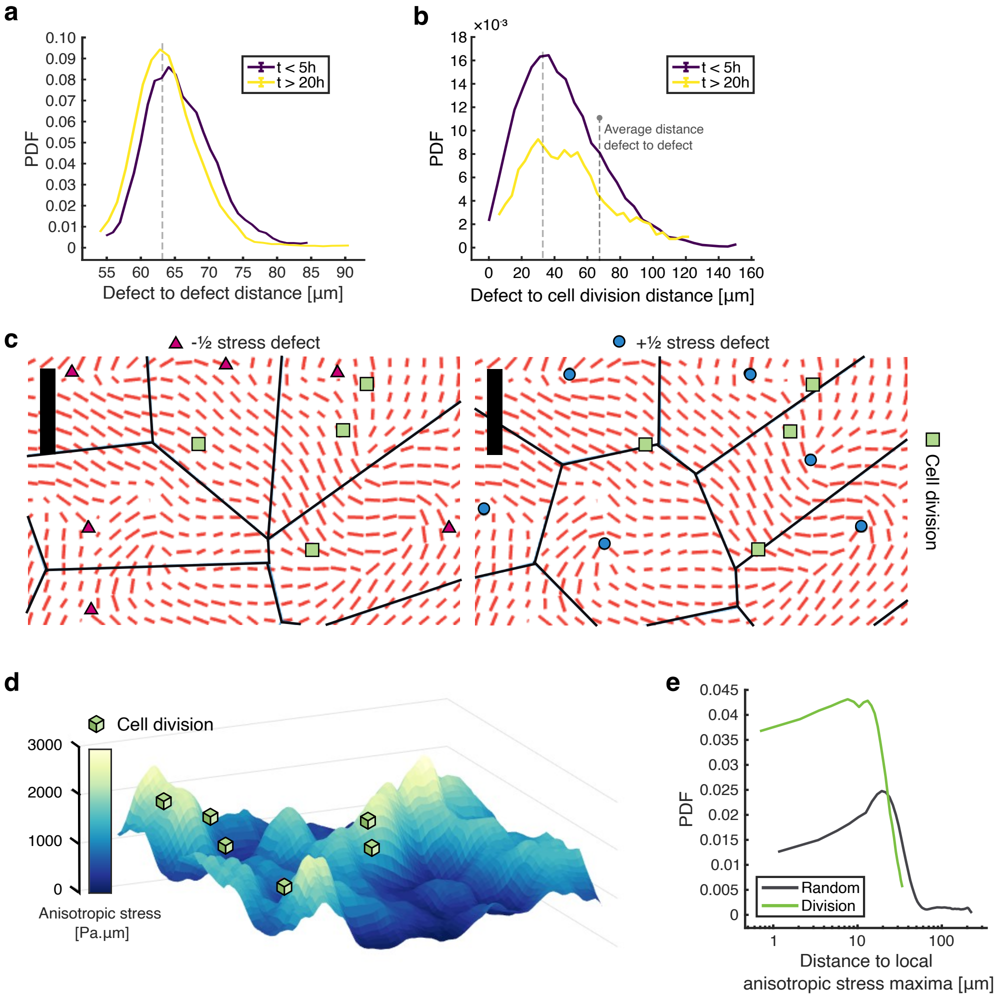


**Fig. S8 Cell division events are localized away from stress topological defects**. **a**) Probability density function (PDF) of average defect to defect distance in one given time (distances are computed on defects with identical charges) for low density and high density. 780 points (low density), 780 points (high density). n = 13 movies from N = 3 independent experiments. **b**) PDF of stress defects to cell division event distance regardless of the charge for liquid and jammed state. 676 divisions (low density), 270 divisions (high density). n = 13 movies from N = 3 independent experiments. **c**) Left, Voronoi tesselation using $-\frac{1}{2}$ stress defects (pink triangle) as generation cores. Right, Voronoi tesselation using $+\frac{1}{2}$ stress defects (blue circle) as generation cores. Principle stress orientation in red. Cell divisions in green square. The selected snapshot is representative of the typical situation during tissue densification (t = 5h). **d**) Mechanical anisotropic levels snapshot in the monolayer (t = 5h), represented as a mechanical potential landscape. Cell divisions (green square) are happening near local maxima. Color code is the anisotropic stress magnitude. **e**) PDF of distance to nearest anisotropic stress maximal for cell divisions and randomly generated points in space and time. 1672 divisions, and 2219 randomly generated points. n = 13 movies from N = 3 independent experiments. [Scale bar 50 µm (c).]

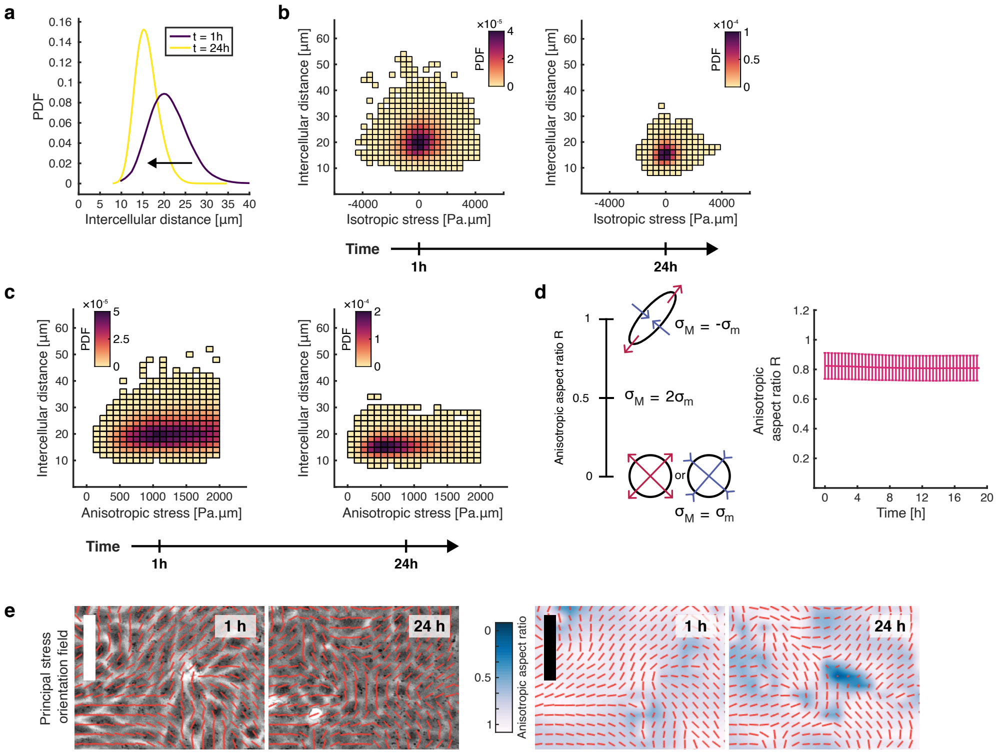


**Fig. S9 Cell density and intercellular stress organization are coupled during tissue densification.** **a**) Probability density function (PDF) of the distribution of intercellular distance between cells in the tissue in low density state (t = 1h) and high density state (t = 24h). n = 15 movies from N = 3 independent experiments. **b**) 2D histogram of the isotropic stress and intercellular distance coupling within MDCK monolayer, left, low cell density (t = 1h), and right, high cell density (t = 24h). Color code is the PDF. n = 15 movies from N = 3 independent experiments. **c**) 2D histogram of the anisotropic stress and intercellular distance coupling within MDCK monolayer, left, low cell density (t = 1h), and right, high cell density (t = 24h). Color code is the PDF. n = 15 movies from N = 3 independent experiments. **d**) Right, simplified representation of the coupling between maximum and minimum stress for various value of anisotropic aspect ratio. Left, average evolution in time of the anisotropic aspect ratio within the whole tissue, for MDCK WT. n = 24 movies from N = 5 independent experiments. **e**) Right, phase contrast image of sub-confluent MDCK monolayer (t = 1h) at low density, and at high density (t = 24h). Left, corresponding anisotropic aspect ratio field for low density and high density. Color code is the anisotropic aspect ratio value. Principal stress orientation is in red. [Scale bar 70 μm (e). All error bars show the confidence interval at 95 %.]

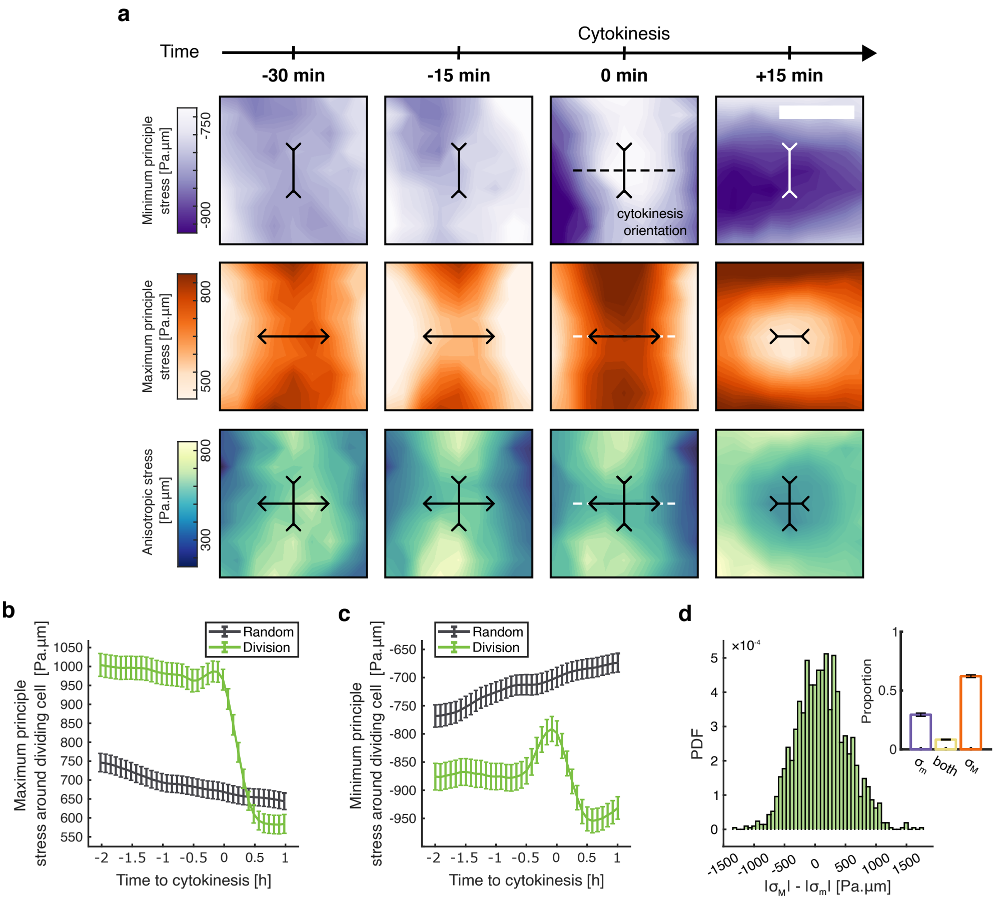


**Fig. S10 The mechanical signature of cytokinesis is anisotropic**. **a**) Stress field around cell division before and after cytokinesis, oriented along the cell division axis (left-right axis). The black double arrow show the orientation of the maximum and minimum stress, with their sign. Top, minimum principle stress, middle, maximum principle stress, and bottom anisotropic stress. 1478 cells division from n = 13 movies from N = 3 independent experiments. **b**) Average maximum principal stress around cell undergoing mitosis. Cytokinesis happens at t = 0h. 1672 cell divisions and 2219 random positions. n = 13 movies from N = 3 independent experiments. **c**) Average minimum principal stress around cell undergoing mitosis. Cytokinesis happens at t = 0h. 1672 cell divisions and 2219 random positions. n = 13 movies from N = 3 independent experiments. **d**) Probability distribution function (PDF) of the absolute contribution of maximum to minimum stress around cell division right before cytokinesis. 1672 cell divisions. Corresponding discrete barplot of the proportion of each contribution possible is also shown. n = 13 movies from N = 3 independent experiments. [Scale bar 25 µm (a). All error bars show the confidence interval at 95 %.]

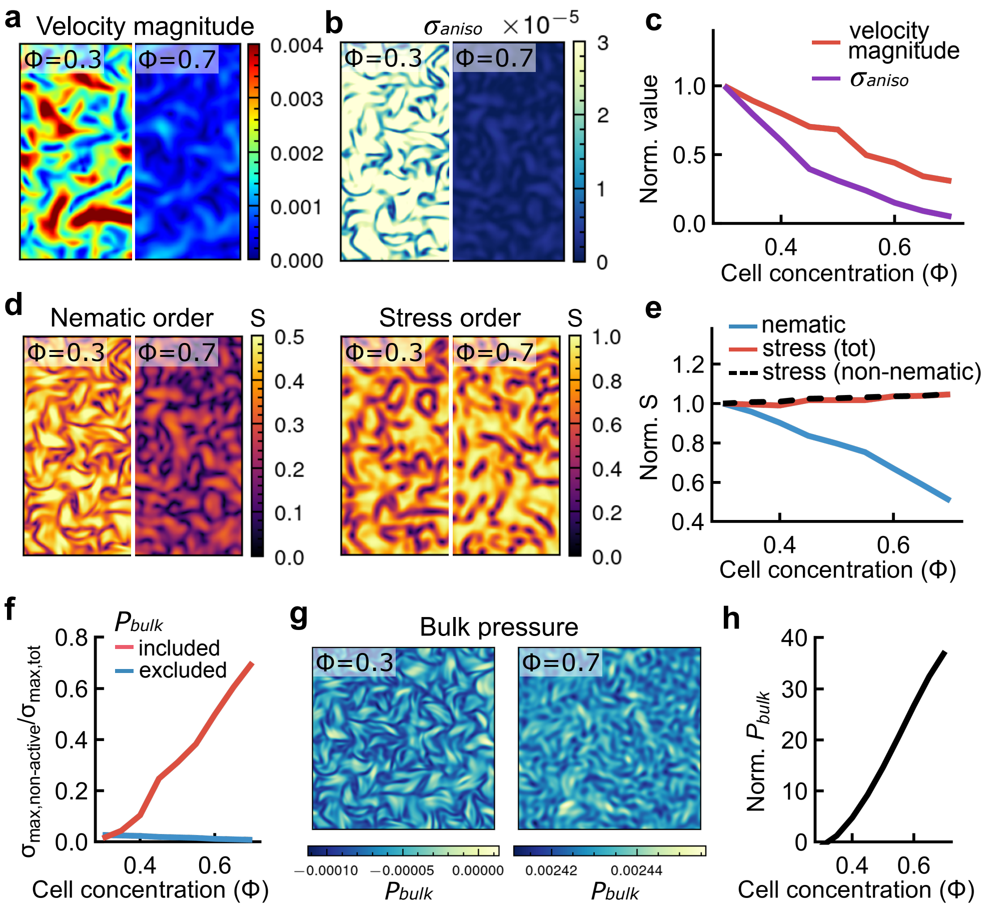


**Fig. S11 Active nematic model with uniformly increased cell density reproduces key features of the tissue jamming transition.** **a**, Velocity magnitude decreases with increasing cell concentration ($\phi$). **b**, Anisotropic stress ($\sigma_{\text{aniso}}$) is progressively reduced at higher $\phi$. **c**, Normalized velocity magnitude and anisotropic stress as functions of $\phi$, both referenced to their respective values at $\phi = 0.3$, showing a coordinated decline with increasing density. **d**, Left: Nematic order collapses as $\phi$ increases. Right: In contrast, principal stress orientation remains robust. **e**, Normalized scalar order parameters for the nematic director (blue), principal stress (red), and principal stress calculated from the stress tensor with nematic stress excluded (black dashed), each referenced to their values at $\phi = 0.3$. While nematic order diminishes, stress and non-nematic stress alignment persists. **f**, Relative contribution of non-nematic stress to total stress, computed as the ratio of the maximal principal stress of non-nematic components ($\sigma_{\text{max,non-nematic}}$) to that of total stress ($\sigma_{\text{max,tot}}$), increases with $\phi$ (red). This trend disappears when bulk pressure ($P_{\text{bulk}}$) is excluded from the non-nematic stress (blue). **g,h)** Bulk pressure ($P_{\text{bulk}}$) rises with increasing cell concentration. [All simulations: $N = 3$; curves represent mean values.]

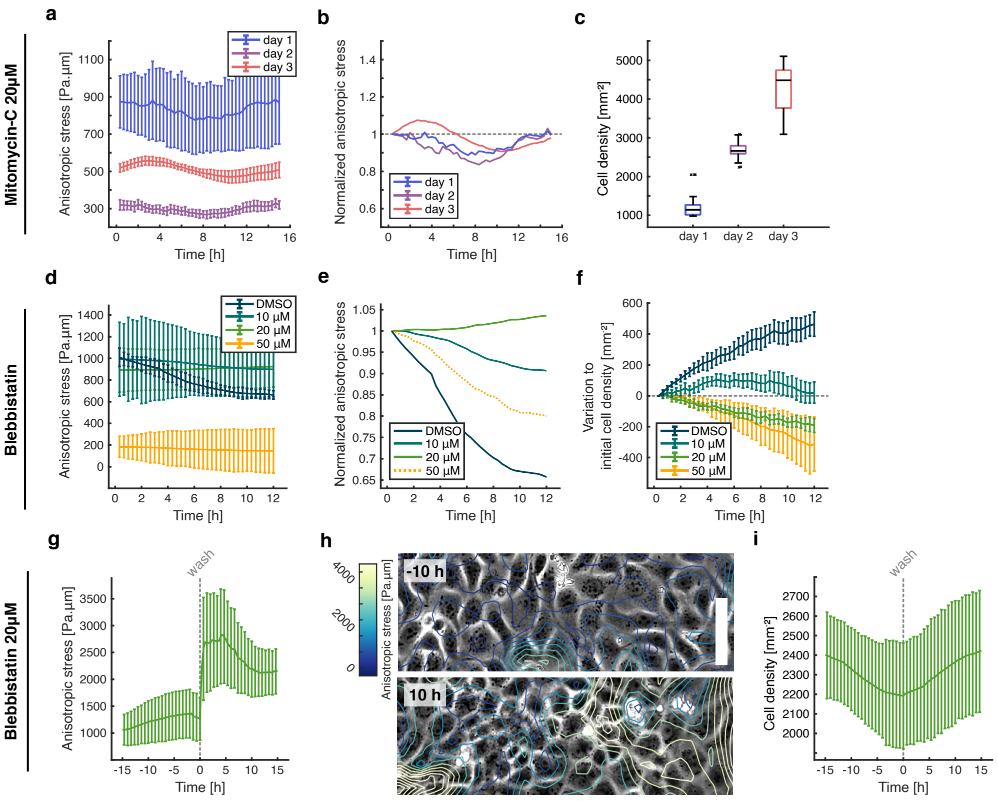


**Fig. S12 Preventing cytokinesis freezes the anisotropic mechanical feedback loop**. **a**) Average (mean) anisotropic stress evolution for confluent MDCK monolayers under mitomycin-C added at various time steps after confluency was reached. n = 14 movies (added a day 1), n = 12 movies (added at day 2), n = 14 movies (added at day 3) from N = 3 independent experiments (day 1) and N = 2 independent experiments (day 2 & day 3). **b**) Average (mean) normalized anisotropic stress evolution, corresponding to g). **c**) Average cell density for each mitomycin conditions, corresponding to g). **d**) Average (mean) anisotropic stress evolution for confluent MDCK monolayers under various doses of blebbistatin, with DMSO as a control. n = 10 movies (10 $\mu$M), n = 10 movies (20 $\mu$M), n = 7 movies (50 $\mu$M), and n = 9 movies (DMSO) from N = 2 independent experiments. **e**) Average (mean) normalized anisotropic stress evolution, corresponding to d). **f**) Average (mean) cell density augmentation regarding the initial cell density in each experiments, corresponding to d). **g**) Average (mean) anisotropic stress evolution for a confluent MDCK monolayer under 20 $\mu$M blebbistatin. A wash of the drug is done at t = 0h. n = 11 movies from N = 2 independent experiments. **h**) Phase contrast image of confluent MDCK monolayer, showing in overlap the local anisotropic stress field, before (t = -10h) and after (t = 10h) blebbistatin wash. **i**) Average (mean) cell density during the course of the experiment corresponding to a). n = 11 movies from N = 2 independent experiments. [Scale bar 50 µm (h). All error bars show the confidence interval at 95 %.]

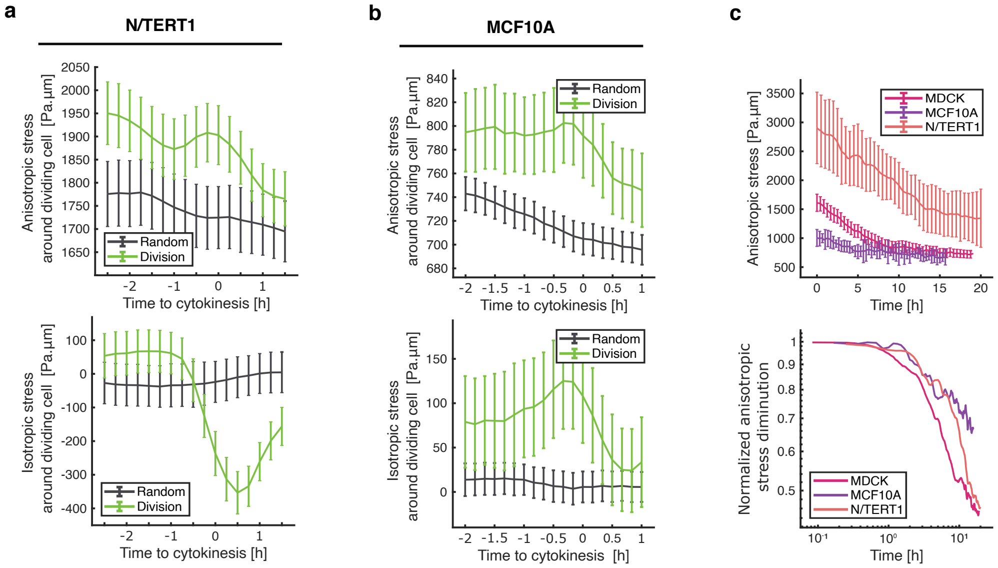


**Fig. S13 Anisotropic stress and cell division feedback loop is recovered in different epithelial cell lines**. **a**) N/TERT1 cells ; top, average anisotropic stress around cells undergoing mitosis, bottom, average isotropic stress around cells undergoing mitosis. Cytokinesis happens at t = 0h. 851 cell divisions and 704 random positions. n = 7 movies from N = 3 independent experiment. **a**) MCF10A cells ; top, average anisotropic stress around cells undergoing mitosis, bottom, average isotropic stress around cells undergoing mitosis. Cytokinesis happens at t = 0h. 354 cell divisions and 2348 random positions. n = 7 movies from N = 2 independent experiments. **c**) Top, average evolution in time of the anisotropic stress during jamming transition for various cell lines, and bottom, corresponding plot normalized by the maximum anisotropic stress levels of each cell types, in a log-log scale. n = 24 movies from N = 5 independent experiments (MDCK), n = 7 movies from N = 2 independent experiments (MCF10A) and n = 7 movies from N = 3 independent experiment (N/TERT1). [All error bars show the confidence interval at 95 %.]

#### Video captions:

**Video S1:** Timelapse video of densification of MDCK WT cells. Frame rate 5 min. Nematic director field in blue. Scale bar 80 µm.

**Video S2:** Timelapse video of densification of MDCK WT cells, with the corresponding isotropic and anisotropic stress field. Frame rate 5 min. Principal stress orientation field in red. Scale bar 80 µm.

**Video S3:** Timelapse video of intestinal organoid-derived 2D monolayer, after three days culture, with villin tomato channel. Principal stress orientation field in red. Frame rate 10 min. Scale bar 80 µm.

**Video S4:** Timelapse video of densification of MDCK WT cells, corresponding to Movie S1. Frame rate 5 min. Principal stress orientation field in red. Scale bar 80 µm.

**Video S5:** Timelapse video of densification of MDCK FUCCI cells. Frame rate 5 min. Magenta is G1/G0 signal, green is S-G2-M signal, overlapped with phase contrast images. Scale bar 80 µm.

**Video S6:** Timelapse video of densification of MCF10A cells, with the corresponding isotropic and anisotropic stress field. Frame rate 10 min. Principal stress orientation field in red. Scale bar 80 µm.

**Video S7:** Timelapse video of densification of N/TERT1 cells, with the corresponding isotropic and anisotropic stress field. Frame rate 15 min. Principal stress orientation field in red. Scale bar 80 µm.
